# DREAM Interrupted: Severing LIN-35-MuvB association in *Caenorhabditis elegans* impairs DREAM function but not its chromatin localization

**DOI:** 10.1101/671024

**Authors:** Paul D. Goetsch, Susan Strome

## Abstract

The mammalian pocket protein family, which includes the Retinoblastoma protein (pRb) and Rb-like pocket proteins p107 and p130, regulates entry into and exit from the cell cycle by repressing cell cycle gene expression. Although pRb plays a dominant role in mammalian systems, p107 and p130 are the ancestral pocket proteins. The Rb-like pocket proteins interact with the highly conserved 5-subunit MuvB complex and an E2F-DP transcription factor heterodimer, forming the DREAM (for <u>D</u>p, <u>R</u>b-like, <u>E</u>2F, <u>a</u>nd <u>M</u>uvB) complex. DREAM complex assembly on chromatin culminates in repression of target genes mediated by the MuvB subcomplex. Here, we examined how the Rb-like pocket protein contributes to DREAM formation and function by disrupting the interaction between the sole *Caenorhabditis elegans* pocket protein LIN-35 and the MuvB subunit LIN-52 using CRISPR/Cas9 targeted mutagenesis. A triple alanine substitution of LIN-52’s LxCxE motif severed LIN-35-MuvB association and caused classical DREAM mutant phenotypes, including synthetic multiple vulvae, high-temperature arrest, and ectopic expression of germline genes in the soma. However, RNA-seq revealed limited upregulation of DREAM target genes when LIN-35-MuvB association was severed, as compared to gene upregulation following LIN-35 loss. Based on chromatin immunoprecipitation, disrupting LIN-35-MuvB association did not affect the chromatin localization of E2F-DP, LIN-35, or MuvB components. In a previous study we showed that in worms lacking LIN-35, E2F-DP and MuvB chromatin occupancy was reduced genome-wide. With LIN-35 present but unable to associate with MuvB, our present study suggests that the E2F-DP-LIN-35 interaction promotes E2F-DP’s chromatin localization, which we hypothesize supports MuvB chromatin occupancy indirectly through DNA. Altogether, this study highlights how the pocket protein’s association with MuvB supports DREAM function but is not required for DREAM’s chromatin occupancy.

## Introduction

Members of the mammalian Retinoblastoma (Rb) protein family, pRb, p107, and p130, collectively called pocket proteins, serve key roles in regulating transcription during the cell cycle (Classon and Dyson 2001; Classon and Harlow 2002; Cobrinik 2005; Burkhart and Sage 2008; Dick and Rubin 2013). In mammalian cells, pRb interacts with activating E2F-DP transcription factor heterodimers (in mammals, E2F1/2/3-DP1/2), sequestering E2F-DP and preventing E2F-DP-mediated activation of early cell cycle genes (Helin *et al*. 1992; Lees *et al*. 1993; Liban *et al*. 2016). In contrast, the Rb-like proteins p107 and p130 interact with repressive E2F-DPs (in mammals, E2F4/5-DP1/2) and a highly conserved 5-subunit MuvB subcomplex (in mammals, LIN9, LIN37, LIN52, LIN54, and RBAP48), forming the 8-subunit DREAM transcriptional repressor complex (Korenjak *et al*. 2004; Lewis *et al*. 2004; Harrison *et al*. 2006; Litovchick *et al*. 2007; Schmit *et al*. 2007). When associated with the DREAM complex, MuvB mediates transcriptional repression of early and late cell cycle genes (Litovchick *et al*. 2007; Goetsch *et al*. 2017; Muller *et al*. 2017). The transcriptional functions of the DREAM complex and pRb overlap, with each being sufficient to establish and maintain cellular quiescence (G0) if the other is inactive (Hurford *et al*. 1997; Litovchick *et al*. 2007; Muller *et al*. 2017). Upon progression into the cell cycle, pRb and the Rb-like pocket proteins are phosphorylated by CDK4/6-cyclin D, releasing their respective interaction partners and triggering activation of cell cycle genes (Tedesco *et al*. 2002; Pilkinton *et al*. 2007; Burke *et al*. 2010). Thus, the association and dissociation of pocket proteins from their respective transcriptional complexes governs the switch between cell cycle quiescence and cell cycle progression.

The Rb-like homologs p130 and p107 are the ancestral pocket proteins and likely the conserved components that mediate cell cycle control among eukaryotes (Cao *et al*. 2010; Liban *et al*. 2017). In *C. elegans*, LIN-35 is the sole pocket protein, most closely resembling p130/p107 (Lu and Horvitz 1998). The pocket protein-associated complex MuvB was isolated in *Drosophila melanogaster* (Korenjak *et al*. 2004; Lewis *et al*. 2004) and *Caenorhabditis elegans* (Harrison *et al*. 2006) before homologs were identified in mammals (Litovchick *et al*. 2007; Pilkinton *et al*. 2007; Schmit *et al*. 2007). The *C. elegans* complex, variably called DRM and DREAM, regulates cell cycle genes and requires MuvB to mediate gene repression (Boxem and van den Heuvel 2002; Goetsch *et al*. 2017). *C. elegans* DREAM also regulates cell-fate specification by antagonizing Ras signaling during vulval development (Myers and Greenwald 2005; Cui *et al*. 2006; Harrison *et al*. 2006) and by protecting somatic cells from expressing germline genes (Wang *et al*. 2005; Petrella *et al*. 2011).

Extensive biochemical analyses have demonstrated how the DREAM complex assembles on chromatin (Figure 1A) (Litovchick *et al*. 2007; Pilkinton *et al*. 2007; Schmit *et al*. 2007; Guiley *et al*. 2015; Asthana *et al*. 2022). E2F-DP and LIN54, a MuvB component, direct site-specific chromatin localization (Zwicker *et al*. 1995; Schmit *et al*. 2009; Muller and Engeland 2010; Muller *et al*. 2012; Marceau *et al*. 2016). The Rb-like pocket protein serves as a bridge between the 2 DNA-binding DREAM components (Guiley *et al*. 2015). LIN52 interacts with the pocket protein via an “LxCxE motif” in LIN52. In mammals, the LxCxE motif is instead a suboptimal LxSxExL sequence that is rendered optimal by phosphorylation of a nearby serine residue (S28) (Guiley *et al*. 2015) (Figure 1B). S28 phosphorylation by DYRK1A kinase induces formation of mammalian DREAM (Litovchick *et al*. 2011). In *C. elegans*, the conserved *lin-52* gene encodes the optimal LxCxE sequence (Figure 1B). *C. elegans* lacks a DYRK1A homolog and its corresponding consensus motif RxSP in LIN-52 (Figure 1B), suggesting that in *C. elegans* a phospho-switch does not induce DREAM formation (Litovchick *et al*. 2011; Guiley *et al*. 2015). Importantly, the LxCxE binding motif mediates a high-affinity interaction that is employed by the human papillomavirus (HPV) viral oncoprotein E7 to prevent association of LIN52 with the mammalian pocket protein p130 (Guiley *et al*. 2015).

**Figure 1.**
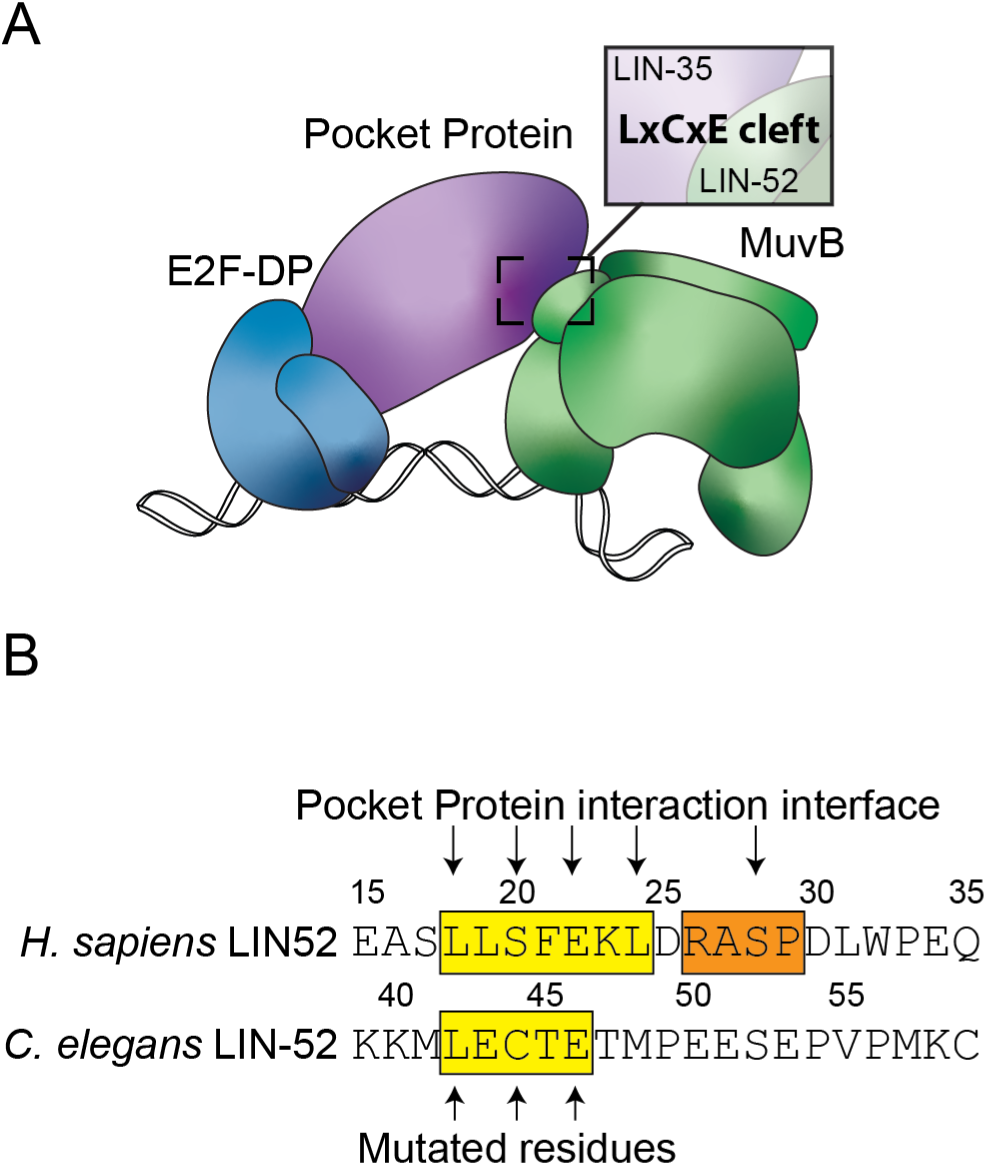
LIN-35 and MuvB associate via the LxCxE motif of LIN-52. (A) Model of the *C. elegans* DREAM complex bound to DNA: E2F-DP (blue), the pocket protein LIN-35 (purple), and the 5-subunit MuvB subcomplex (green). The highlighted region shows the target region for this study: an LxCxE binding motif in the MuvB subunit LIN-52 that interacts directly with the LIN-35 pocket protein. (B) Alignment of *H. sapiens* LIN52 and *C. elegans* LIN-52 sequences. The human LxSxExL and worm LxCxE sequences are highlighted in yellow, and the human DYRK1A consensus phosphorylation sequence is highlighted in orange. Arrows indicate residues involved in the interaction with the pocket protein and amino acid residues converted to alanine in this study.

Here, we assessed how the Rb-like pocket protein contributes to DREAM complex formation and function on chromatin. We previously reported that the absence of LIN-35 results in a genome-wide decrease in chromatin occupancy of both E2F-DP and MuvB, illustrating how DRM/DREAM disassembly likely proceeds during cell cycle progression (Goetsch *et al*. 2017). The model of DREAM complex assembly centers on reintroduction of the pocket protein associations with E2F-DP and MuvB as cells finish the cell cycle (Guiley *et al*. 2015). To test this model, we used CRISPR/Cas9-mediated genome editing of the *C. elegans* LIN-52 subunit of MuvB to sever the association of LIN-35 with the MuvB subcomplex. Disrupting that association caused phenotypes associated with impairment of DREAM function, including a highly penetrant synthetic multivulval (SynMuv) phenotype (Fay and Yochem 2007), high-temperature larval arrest (Petrella *et al*. 2011), and ectopic expression of germline genes in the soma (Wang *et al*. 2005; Petrella *et al*. 2011). However, our genome-wide transcript analysis revealed that severing the association of LIN-35 and LIN-52 led to upregulation of a relatively small set of DREAM target genes, which displayed a lower magnitude of upregulation than caused by loss of LIN-35. Moreover, chromatin immunoprecipitation (ChIP) revealed that the chromatin association of E2F-DP-LIN-35 and MuvB was not impaired by loss of LIN-35-MuvB association, even in gene promoters of upregulated DREAM target genes. Altogether, our results indicate that loss of the direct association between LIN-35 and MuvB causes limited upregulation of DREAM target genes, likely due to DREAM chromatin occupancy being relatively normal, but even those minimal effects lead to phenotypes consistent with impaired DREAM function in *C. elegans*.

## Materials and Methods

### Worm strains

Strains were cultured on Nematode Growth Medium (NGM) agarose plates with *E. coli* OP50 and incubated at 20°C. The following strains were used: Wild type N2 (Bristol), SS1240 *lin-52(bn132(lin-52p::TagRFP-T^SEC^3xFLAG::lin-52 3’ UTR)) III / hT2G [bli-4(e937) let-?(q782) qIs48] (I:III)*, SS1241 *lin-52(bn133(lin-52p::TagRFP-T::3xFLAG::lin-52 3’ UTR)) III / hT2G [bli-4(e937) let-?(q782) qIs48] (I:III)*, SS1325 *lin-52(bn138(lin-52::GFP^SEC^3xFLAG)) III*, SS1256 *lin-52(bn139(lin-52::GFP::3xFLAG)) III*, SS1273 *lin-52(bn150(lin-52[C44A]::GFP::3xFLAG)) III*, SS1276 *lin-52(bn151(lin-52[L42A,C44A,E46A]::GFP ::3xFLAG)) III*, SS1350 *lin-52(bn139) III; lin-15A(n767) X*, SS1351 *lin-52(bn150) III; lin-15A(n767) X*, SS1352 *lin-52(bn151) III; lin-15A(n767) X*, SS1406 *lin-8(n2731) II; lin-52(bn139) III*, SS1407 *lin-8(n2731) II; lin-52(bn150) III*, SS1408 *lin-8(n2731) II; lin-52(bn151) III*, PDG29 *lin-52(bn139) III; pgl-1(sam52[pgl-1::mTagRFPT::3xFLAG]) IV*, PDG30 *lin-52(bn150) III; pgl-1(sam52) IV*, and PDG31 *lin-52(bn151) III*; *pgl-1(sam52) IV*.

### CRISPR/Cas9-mediated genome editing

For all genomic edits, 20 nucleotide crDNA targeting sequences were identified using the MIT CRISPR design tool (http://crispr.mit.edu). Single guide RNA sequences were cloned into the PU6::unc119_sgRNA vector (Addgene plasmid #46169) using the overlapping PCR fragment method described in (Friedland *et al*. 2013) or were cloned into pDD162 (Addgene plasmid #47549) using the Q5 Site Directed Mutagenesis Kit (New England Biolabs), as described in (Dickinson *et al*. 2013). Homologous repair templates were cloned into pDD282 (Addgene plasmid #66823) or pDD284 (Addgene plasmid #66825) using Glibson Assembly (New England Biolabs) (Gibson *et al*. 2009), as described in (Dickinson *et al*. 2015). CRISPR/Cas9 component plasmids were co-injected with marker plasmids (Frokjaer-Jensen *et al*. 2008) to identify strains with an extra-chromosomal array instead of a mutated endogenous gene. For targeted mutagenesis, *dpy-10(cn64)* sgRNA (pJA58, Addgene plasmid #59933), and *dpy-10(cn64)* ssDNA template, *dpy-10(cn64)* guide and ssDNA template were co-injected to select for positive CRISPR activity in injectant progeny, as described in (Arribere *et al*. 2014). Additional details are provided in Supplemental Materials and Methods.

### Microscopy

L1 and L4 larvae were mounted on a 10% agarose pad and immobilized in a 1-2 μL suspension of 0.1 μm polystyrene beads (Polysciences), as described in (Kim *et al*. 2013). Fluorescence images of L1 larvae were acquired using a Leica DM4B upright microscope with QImaging QIClick camera. Fluorescence images of L4 larvae were acquired using a Solamere spinning-disk confocal system with μManager software (Edelstein *et al*. 2014). The confocal microscope setup was as follows: Yokogawa CSUX-1 spinning disk scanner, Nikon TE2000-E inverted stand, Hamamatsu ImageEM X2 camera, solid state 405-, 488-, and 561-nm laser lines, 435–485, 500–550, and 573–613 fluorescent filters, and Nikon Plan Fluor 40x air objective. Images were processed using Image J (Schneider *et al*. 2012).

### *C. elegans* phenotype scoring

For brood size analyses, L4 individuals were cloned to fresh plates every 24 hours and all progeny were counted. For SynMuv phenotype scoring, 3 replicate plates per strain were set up with 5-10 adults that were allowed to lay eggs for 6 hours. Progeny were incubated at 20°C for 3 days, then scored for the presence or absence of pseudovulvae. The percentages of multivulva worms in each replicate population were averaged, and the standard deviation was calculated. For HTA phenotype scoring, L4 larvae of each strain were incubated at 24°C or 26°C overnight, then 3 plates per strain were set up with 9-12 adults that were allowed to lay eggs for 4-5 hours. Progeny incubated at 24°C or 26°C were counted 2 days later and scored as arrested larvae (HTA) or adults (not HTA). Results from each plate were combined for each strain.

### Immunoblotting and co-immunoprecipitation (co-IP)

For immunoblotting whole worm lysates, 200 adults from each strain were picked into SDS gel-loading buffer (50 mM pH 6.8 Tris-Cl, 2% sodium dodecyl sulfate, 0.1% bromophenol blue, 100 mM β-mercaptoethanol). For co-IP, embryos collected after bleaching gravid worms were aged for 3.5 hours and then frozen in liquid nitrogen, and lysates were prepared as described in (Goetsch *et al*. 2017). For each IP, 8 mg of protein lysate was mixed with antibody-conjugated Dynabeads (ThermoFisher) and incubated for 2 hours at 4°C. Proteins were separated by SDS/PAGE, and western blot analysis was performed using a 1:1,000-1:5000 dilution of primary antibody and 1:2,000 dilution of an appropriate HRP-conjugated secondary antibody. Serial western blot analysis was performed by stripping the blot with buffer containing 0.2M pH 2.2 glycine, 0.1% SDS, and 1% Tween-20 between antibody probings. Additional details are provided in Supplemental Materials and Methods.

### Chromatin immunoprecipitation (ChIP) and sequential ChIP

Embryos collected after bleaching gravid worms were aged for 3.5 hours and then frozen in liquid nitrogen. Lysates were prepared by grinding, crosslinking for 10 minutes in 1% formaldehyde, and sonicating to an average size of 250 base pairs in FA buffer (50 mM HEPES/KOH pH 7.5, 1 mM EDTA, 1% Triton X-100, 0.1% sodium deoxycholate, 150 mM NaCl) using a Bioruptor (Diagenode) on the high setting with 60 rounds of 30 seconds on and 1 minute rest. Protein concentrations of lysates were determined using a Qubit fluorometer. ChIP and sequential ChIP experiments were performed as described in (Goetsch *et al*. 2017) and in Supplemental Materials and Methods. Quantitative PCR was performed using SYBR green reagents on an Applied Biosystems QuantStudio 3 and ViiA 7 Real-Time PCR Systems (ThermoFisher).

### Analysis of transcript levels by RT-qPCR and RNA-seq

Embryos collected after bleaching gravid worms were aged for 3.5 hours and then frozen in Trizol for RNA isolation. A total of 1 μg RNA was treated with DNase and reverse transcribed using the High Capacity cDNA Kit (Applied Biosystems). qPCR was performed using SYBR green reagents on an Applied Biosystems QuantStudio 3 Real-Time PCR System (ThermoFisher). The relative quantity of experimental transcripts was calculated with *act-2* as the control gene using the ΔCt method.

RNA-seq libraries were prepared using the NEBNext Poly(A) mRNA Magnetic Isolation Module and NEBNext Ultra II Directional RNA Library Prep Kit for Illumina (New England Biolabs). Samples were multiplexed using the NEBNext Multiplex Oligos for Illumina (NEB). Libraries were sequenced on an Illumina HiSeq X system by Novogene Corporation Inc. (Davis, CA) to acquire 150-bp paired-end reads. Sequence reads were trimmed using fastp (Chen *et al*. 2018) and mapped to transcriptome version WBcel235 using STAR (Dobin *et al*. 2013). Read counts per transcript were obtained using HTSeq (Anders *et al*. 2015) and differentially expressed genes were assessed using DEseq2 (Love *et al*. 2014).

### Quantification and statistical analysis

For brood size analysis, significance was determined using a Wilcoxon-Mann-Whitney test comparing CRISPR/Cas9-genome edited strains to wild type (N2). For genome-wide differential expression analysis of *lin-52(3A)* versus *lin-52(WT)* late embryos, we used a 1.5-fold change and adjusted p-value less than 0.05 cutoff as calculated by the Benjamini-Hochberg method performed by DEseq2. For expression analysis of *lin-35(n745)* vs N2 L1s, statistical analysis was performed using R using the Quantile normalization and Robust Multichip Average (RMA) algorithm in the affy package of Bioconductor using the a 1.5-fold change and adjusted p-value less than 0.05 cutoff (Bolstad *et al*. 2003; Irizarry *et al*. 2003). For transcript level analysis by RT-qPCR, significance was determined using a student’s t-test between *lin-35* versus wild type (N2) or *lin-52(1A) or lin-52(3A)* versus *lin-52(WT)*. For ChIP-Qpcr experiments assessing individual subunit’s occupancy changes between strains, significance was determined using a student’s t-test between *lin-35* versus wild type (N2) or *lin-52(3A)* versus *lin-52(WT)*.

### Data and reagent availability

Requests for information, strains, and reagents should be directed to and will be fulfilled by Paul D. Goetsch. Supplemental material is available at FigShare. Supplemental information includes supplemental Materials and Methods. Figure S1 contains full western blots presented in Figure 2D. Figure S2 contains full western blots presented in Figure 3B. Figure S3 contains additional DREAM target genes tested using qPCR for ChIP and sequential ChIP analyses to complement Figure 6. Antibodies used for western blot and ChIP analyses are available in Supplemental Table S1. Primers used for cloning and qPCR are available in Supplemental Table S2. DESeq2 differential gene expression analysis data are available in Supplemental Table S3. RNA-seq expression data used in this study are available from the Gene Expression Omnibus (GEO; http://www.ncbi.nlm.nih.gov/geo) through GEO Series accession number GSE199287. Microarray expression analysis data used in this study are available through GEO Series accession number GSE6547 (Kirienko and Fay 2007).

**Figure 2.**
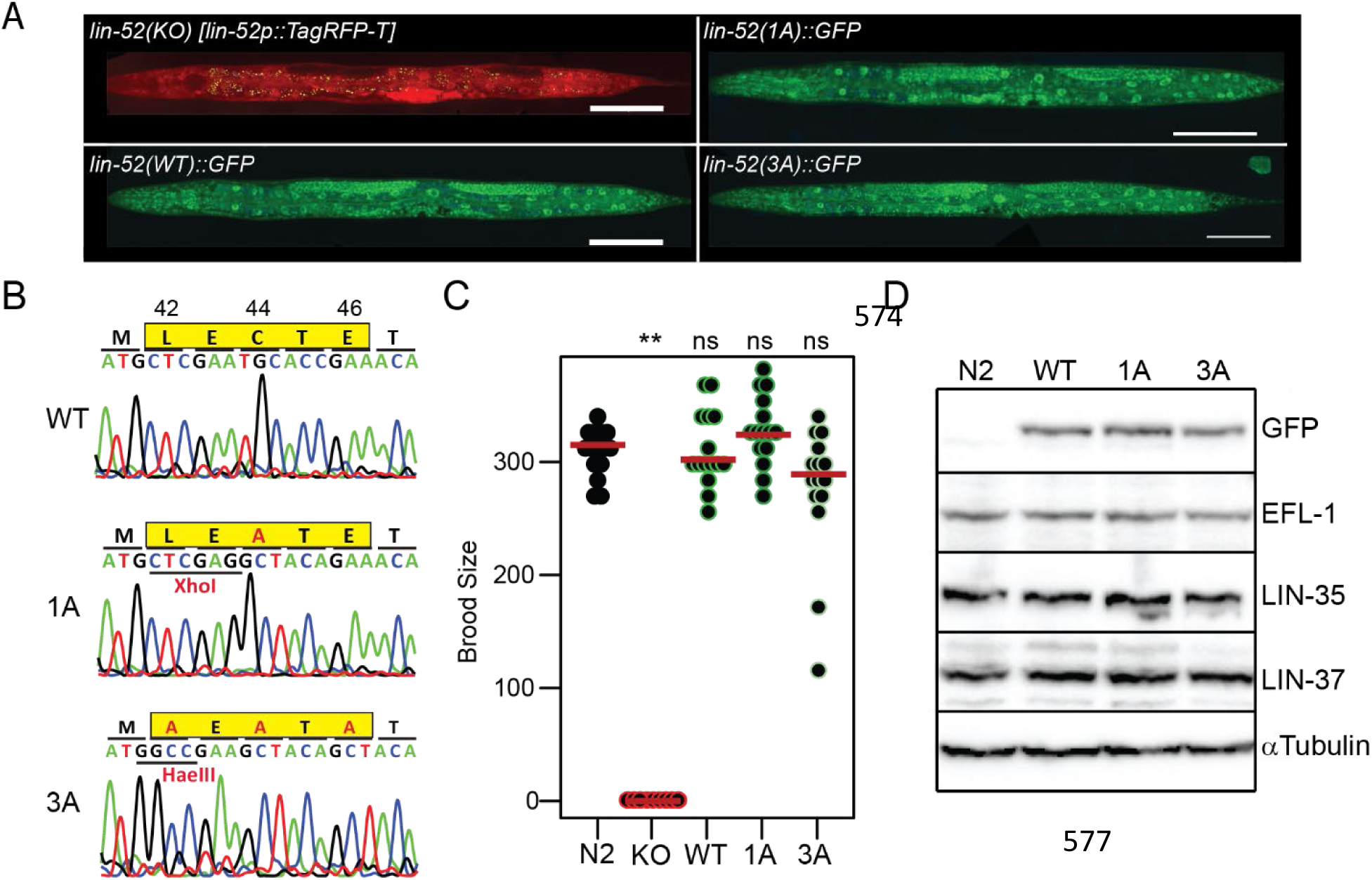
Targeted mutagenesis to disrupt DREAM complex formation. (A) Live worm fluorescence images of *lin-52(KO), lin-52(WT*), *lin-52(1A)*, and *lin-52(3A)* L4 larvae. Composites were artificially straightened. Scale bars, 100μM. (B) Sanger sequencing of the *lin-52* LxCxE coding region (highlighted in yellow) in *lin-52(WT), lin-52(1A*), and *lin-52(3A*). (C) Swarm plots of the brood sizes of wild-type (N2) worms and *lin-52(KO), lin-52(WT), lin-52(1A)*, and *lin-52(3A)* transgenic worms. Significance (** p-value < 0.01) was determined by a Wilcoxon-Mann-Whitney test comparing the indicated strains to wild type (N2). (D) Western blot analysis of DREAM subunits LIN-52 (via GFP tag), EFL-1, LIN-35, and LIN-37 using lysates from wild-type (N2) worms and *lin-52(WT), lin-52(1A)*, and *lin-52(3A)* transgenic worms separated by SDS PAGE. Antibodies used are indicated on the right. Alpha-tubulin was used as a loading control. Full blots are shown in Figure S1.

**Figure 3.**
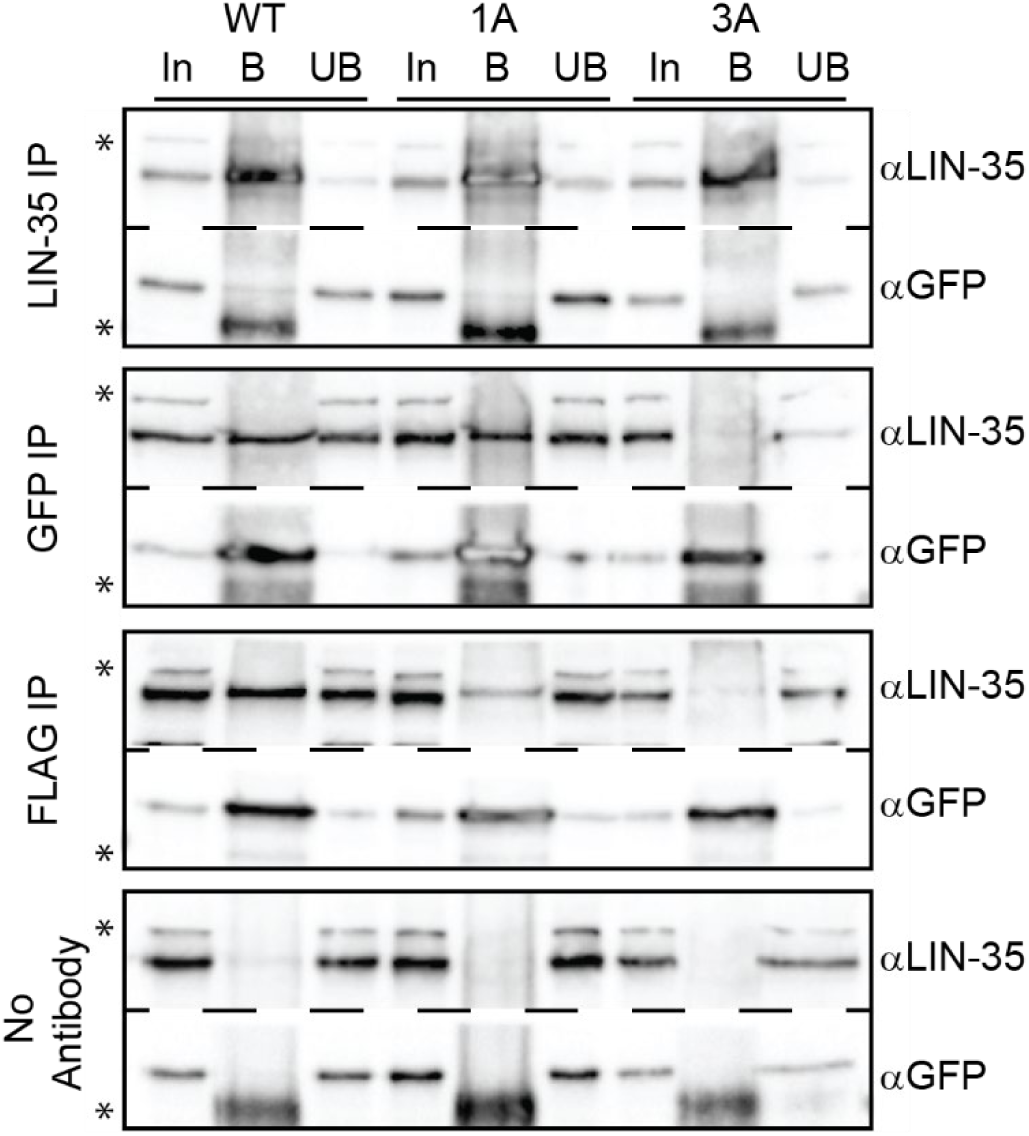
*lin-52* LxCxE binding motif mutants block DREAM formation. Late embryo extracts from *lin-52(WT), lin-52(1A)*, and *lin-52(3A)* (each tagged with GFP and FLAG) were immunoprecipitated with anti-LIN-35, anti-GFP, and anti-FLAG antibodies, with no antibody serving as a negative control. Proteins bound (B) and unbound (UB) were separated by SDS PAGE, and western blot analysis was performed using the antibodies indicated on the right. 5% of Input (In) is shown on the left. Asterisks indicate non-specific bands. Full blots are shown in Figure S2.

## Results

### Targeted mutagenesis to disrupt *C. elegans* DREAM complex formation

Structural studies previously demonstrated that MuvB interacts with the pocket protein via the LIN52 subunit (Figure 1A) (Guiley *et al*. 2015). Using the self-excising cassette (SEC) method for *C. elegans* CRISPR/Cas9 genome editing (Dickinson *et al*. 2015), we generated a *lin-52(KO)* strain (*lin52(bn133[lin-52p::TagRFP-T::3xFLAG]*) by completely replacing *lin-52* coding sequence with *TagRFP-T* coding sequence (Figure 2A). We observed that *lin-52(KO)* rendered worms sterile (Figure 2C), as previously observed in the *lin-52(n3718)* protein null strain (Ceol *et al*. 2006; Harrison *et al*. 2006). This resembles loss of other MuvB components, as loss of LIN-9, LIN-53 (*C. elegans* RBAP48), or LIN-54 in protein null strains also renders worms sterile and affects the levels of other MuvB subunits, suggesting that MuvB components require co-expression for assembly/stability of the complex (Harrison *et al*. 2006). Loss of LIN-37 does not cause sterility and does not affect assembly of the rest of MuvB in either *C. elegans* or mammalian cells (Harrison *et al*. 2006; Mages *et al*. 2017). We next replaced the *TagRFP-T* coding sequence with *lin-52* tagged with a C-terminal *GFP-3xFLAG* coding sequence, generating the *lin-52(WT)* strain (*lin-52(bn139[lin-52::GFP::3xFLAG])*, Figure 2A). *lin-52(WT)* completely rescued fertility (Figure 2C), indicating that the GFP tag does not impair LIN-52 function.

Since LIN-52 is essential for *C. elegans* fertility, we sought to disrupt the LIN-35-LIN-52 interaction without affecting protein integrity. We directed targeted mutagenesis of the LIN-52 LxCxE sequence (Figure 1B) using CRISPR/Cas9-mediated genomic editing. We generated 2 mutants of the LxCxE binding motif in *lin-52(WT)* using the *dpy-10* co-CRISPR method of small oligo homology-directed repair (Arribere *et al*. 2014). We generated the *lin-52(1A)* single-alanine mutation strain (*lin-52(bn150[lin-52[C44A]::GFP::3xFLAG)*) and the *lin-52(3A)* triple-alanine mutation strain (*lin-52(bn151(lin-52[L42A,C44A,E46A]::GFP::3xFLAG)*) (Figure 2B) with the intent to completely disrupt LIN-52’s interaction with the *C. elegans* pocket protein LIN-35. Additional silent mutations were included in the oligo repair templates to generate new restriction enzyme cut sites to aid in genotyping (Figure 2B).

Full loss of *C. elegans* DREAM activity causes sterility, as observed in protein null mutants of worm E2F-DP (*dpl-1* and *efl-1)* and worm MuvB (*lin-9, lin-52, lin-53, and lin-54)* (Beitel *et al*. 2000; Chi and Reinke 2006; Tabuchi *et al*. 2011). Since the C-terminally GFP-tagged *lin-52* coding sequence completely rescued *lin-52(KO)* sterility, we were able to test whether *lin-52(1A)* and *lin-52(3A)* impair DREAM function. We observed that neither the 1A nor 3A mutation in the LIN-52 LxCxE sequence caused a significant reduction in brood size (Figure 2C). Using western blot analysis of selected DREAM components from *lin-52(WT)* and mutant lysates, we observed that DREAM component protein levels were unaffected compared to wild type (N2) (Figure 2D, Figure S1). Similarly, using live-image analysis of *lin-52(WT), lin-52(1A)*, and *lin-52(3A)* L4 larvae, we observed that LIN-52 level and localization appeared normal in mutants (Figure 2A). Together, these results demonstrate that mutation of the LIN-52 LxCxE sequence does not cause a *lin-52* null phenotype and does not alter the levels and tissue distribution of MuvB components.

### The 3A substitution in the LIN-52 LxCxE sequence blocks DREAM assembly

To test whether our CRISPR/Cas9-generated 1A and 3A substitutions disrupt the LIN-35-MuvB association, we performed co-immunoprecipitations (co-IPs) from protein extracts prepared from *lin-52(WT), lin-52(1A)*, and *lin-52(3A)* late embryos. We pulled down LIN-35 and tested for LIN-52 association using the GFP epitope, and we pulled down LIN-52 using either the GFP or FLAG epitope and tested for LIN-35 association (Figure 3, Figure S2). In both co-IP experiments, we observed that LIN-52 association with LIN-35 was lost in *lin-52(3A)* extracts but not in *lin-52(1A)* extracts. In 1A extracts, we observed a weaker association between LIN-52 and LIN-35 when pulling down with an anti-FLAG antibody, but the effect was not replicated with the anti-GFP antibody.

The inconsistent results suggest that the 1A substitution weakens but does not eliminate the LIN-52-LIN-35 interaction. Altogether, these results demonstrate that a 3A substitution in the LxCxE sequence is required to sever the protein-protein association between LIN-52 and LIN-35 in *C. elegans*.

### Blocking DREAM complex formation causes classic DREAM loss-of-function phenotypes

DREAM loss-of-function mutations when paired with SynMuv A gene mutations cause a synthetic multivulval (SynMuv) phenotype (Fay and Yochem 2007), high-temperature larval arrest (Petrella *et al*. 2011), and ectopic expression of germline genes in the soma (Wang *et al*. 2005; Petrella *et al*. 2011). We reasoned that if DREAM function was affected by severing the LIN-52-LIN-35 association, then we should observe each of the 3 phenotypes above. When paired with SynMuv A allele *lin-8(n2731)* (Harrison *et al*. 2007) or *lin-15A(n767)* (Huang *et al*. 1994), *lin-52(3A)* resulted in a SynMuv phenotype; *lin-52(WT)* and *lin-52(1A)* did not have this effect (Figure 4A). When worms were grown at high temperature (26°C), *lin-35(3A)* displayed larval arrest; *lin-52(WT)* and *lin-52(1A)* did not cause larval arrest (Figure 4B). When RT-qPCR was used to examine ectopic expression of the germline genes *pgl-1, pgl-3*, and *glh-1, lin-52(3A)* caused upregulation of all 3 genes; *lin-52(1A)* caused upregulaton of *pgl-1* and *pgl-3* but not *glh-1* (Figure 4C). To examine ectopic expression by another method, we crossed a germline-specific *pgl-1::rfp* reporter (Marnik *et al*. 2019) into our *lin-52* mutant strains. We observed ectopic expression of *pgl-1::rfp* in *lin-52(1A)* and *lin-52(3A)* in the intestinal cells of larvae (Figure 4D, E), as expected (Petrella *et al*. 2011). These phenotypes along with our co-immunoprecipitation findings demonstrate that the 3A substitution has a greater impact on DREAM assembly and function than the 1A substitution.

**Figure 4.**
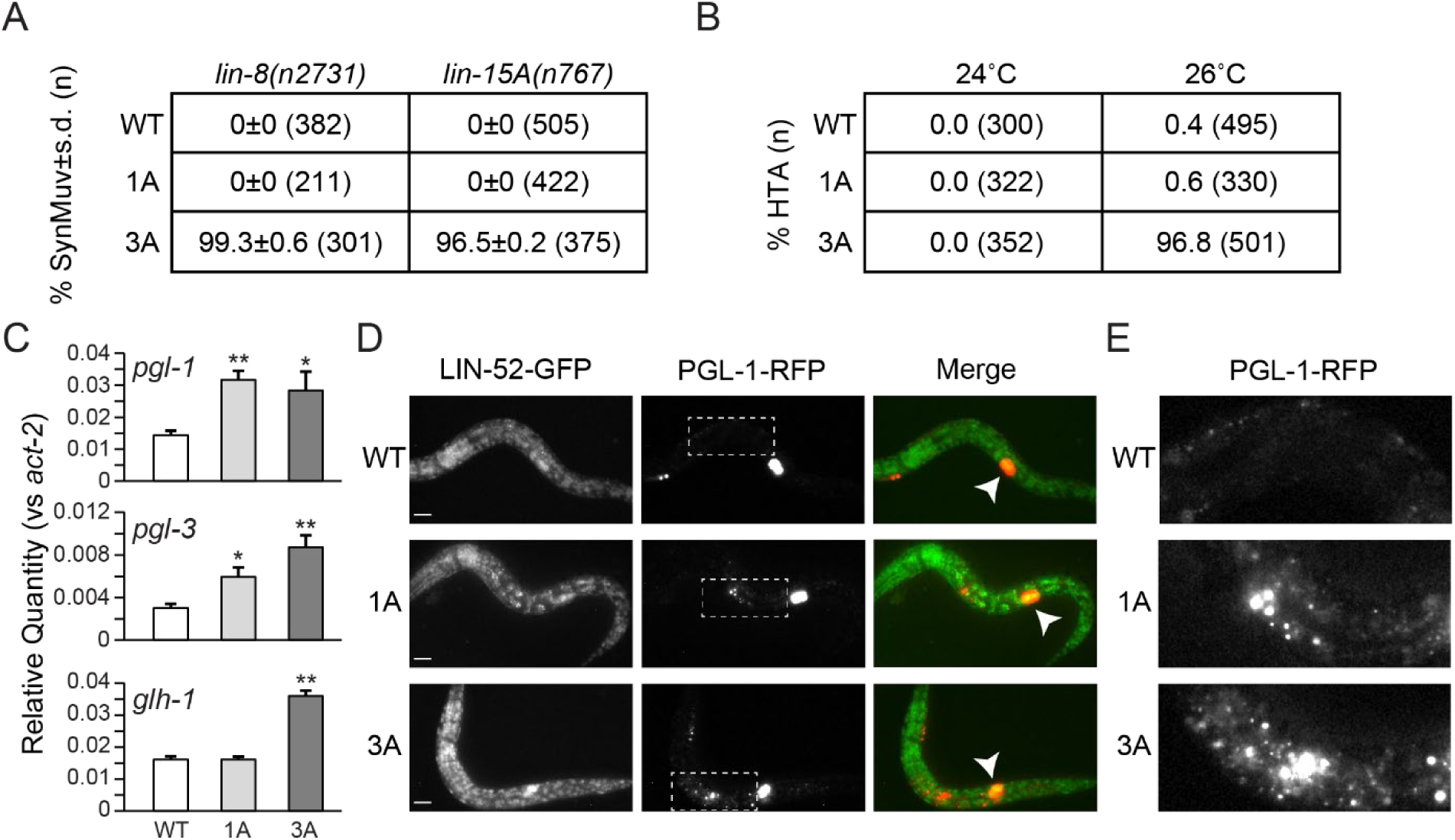
Mutagenesis of the *lin-52* LxCxE binding motif causes SynMuv B phenotypes. (A) Table indicating the percentage Synthetic Multivulval (SynMuv) worms when *lin-52(WT), lin-52(1A)*, and *lin-52(3A)* were combined with SynMuv A mutant alleles *lin-8(n2731)* or *lin-15A(n767)* with standard deviation indicated. The population size (n) is indicated in parentheses. (B) Table indicating the percentage of High Temperature Arrest (HTA) observed in *lin-52(WT), lin-52(1A), and lin-52(3A)* incubated at 24°C and 26°C. The population size (n) is indicated in parentheses. (C) RT-qPCR analysis comparing transcript levels of 3 germline genes (*pgl-1, pgl-3*, and *glh-1*) in *lin-52(WT)* (white), *lin-52(1A)* (light grey), and *lin-52(3A)* (dark grey) late embryos. Expression values from 6 biological replicates were averaged and are presented as the relative quantity compared to *act-2*. Error bars indicate standard error of the mean, and significance was determined by a student’s t-test between transcript levels in mutant (3A or 1A) versus WT (* p-value < 0.05, ** p-value < 0.01). (D,E) Live worm fluorescence images of *lin-52(WT*), *lin-52(1A)*, and *lin-52(3A)* L1 larvae containing a PGL-1::RFP reporter gene. White arrowheads indicate the primordial germ cells Z2 and Z3. Scale bars, 10 μm. The white boxes in (D) indicate regions of ectopic expression of PGL-1::RFP in the intestine of *lin-52(1A)* and *lin-52(3A)* shown in (E).

### Severing the LIN-35-MuvB connection has a minimal effect on repression of DREAM target genes

To investigate whether blocking DREAM assembly affects transcriptional repression of DREAM target genes, we performed whole-genome RNA-sequencing (RNA-seq) in *lin-52(WT)* and *lin-52(3A)* late embryos and compared our data to microarray-based expression data from wild-type and *lin-35(n745)* null mutant L1 larvae (Kirienko and Fay 2007). Our RNA-seq analysis detected 16,855 transcripts (>10 counts across samples), as compared to 16,810 transcripts tested in the microarray experiment (Supplemental Table S3, S4). A total of 13,721 transcripts are shared between the two experiments, including 1,370 of the 1,515 genes previously reported to have DREAM bound to their promoters, which we categorize as DREAM targets (Goetsch *et al*. 2017). In *lin-35(n745)* versus wild-type (N2) L1 larvae, 798 genes (246 DREAM targets) were upregulated and 540 genes (31 DREAM targets) were downregulated (Figure 5A). In contrast, in *lin-52(3A)* versus *lin-52(WT)* late embryos, 102 genes (32 DREAM targets) were upregulated and 9 genes (0 DREAM targets) were downregulated (Figure 5B). Altogether, our transcript analyses reveal that the 3A substitution has a minimal effect on DREAM target regulation, with only ∼2% (32/1370) of DREAM targets upregulated in *lin-52(3A)* late embryos compared to ∼18% (246/1370) of DREAM targets upregulated in *lin-35(n745)* L1s.

**Figure 5.**
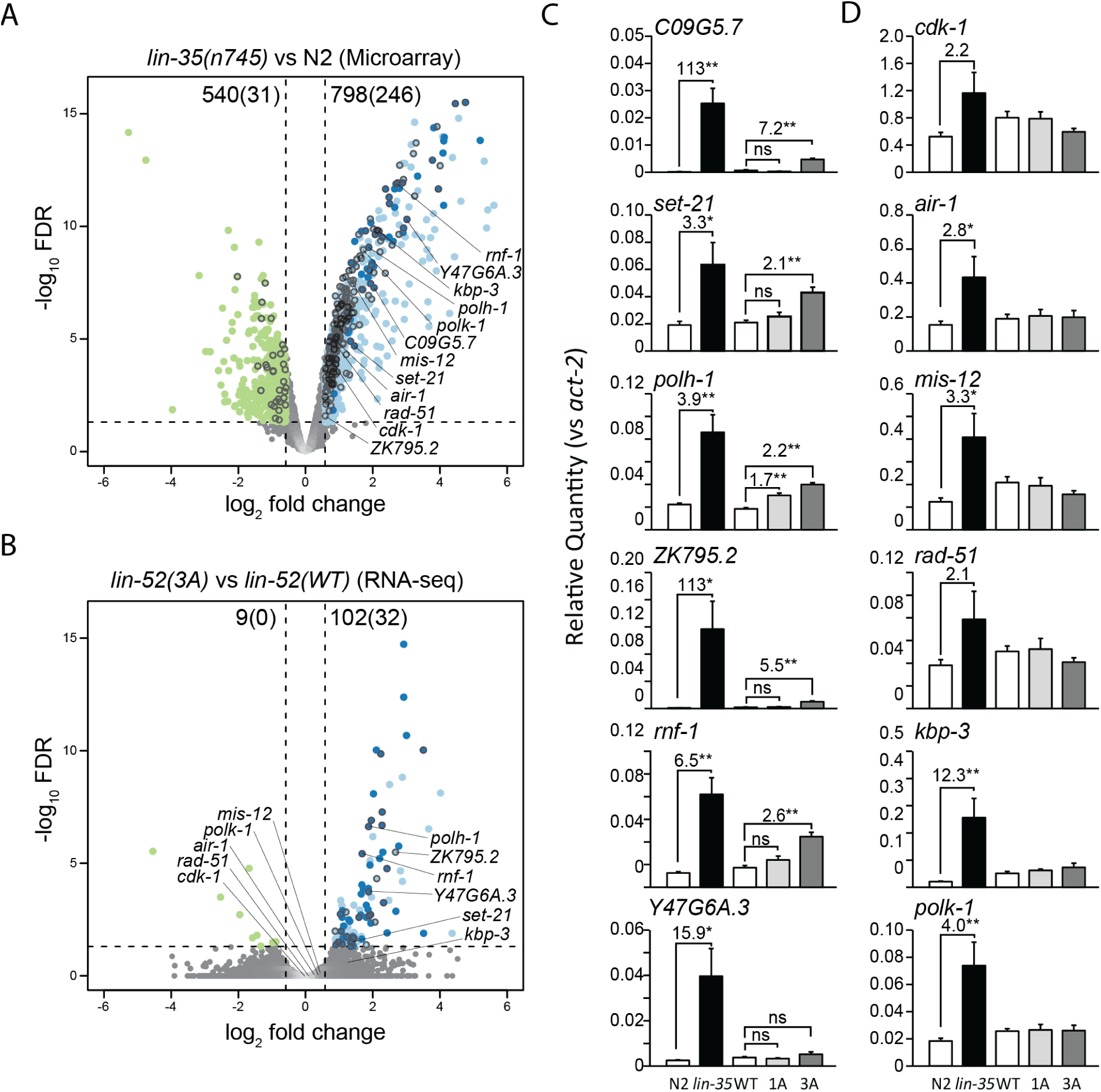
Disruption of DREAM formation leads to upregulation of a small subset of target genes. (A,B) Volcano plots of log2 fold change in transcript levels versus log10 False Discovery Rate (FDR) of 13,721 genes shared between (A) *lin-35(n745)* versus N2 L1 microarray expression analysis reported in (Kirienko and Fay 2007) and (B) *lin-52(3A)* versus *lin-52(WT)* late embryo RNA-seq. Genes downregulated (green) or upregulated (blue) are highlighted, with dark blue indicating upregulated genes observed in both *lin-35* and *lin-52(3A)* data sets. Black circle outlines indicate DREAM target genes, as reported in (Goetsch *et al*. 2017). The number of genes differentially expressed in each analysis are indicated at the top of each plot, with the number of DREAM target genes differentially expressed indicated in parentheses. Dashed lines indicate the significance cut-off of q = 0.05 (horizontal lines) and a 1.5-fold change in transcript level (vertical lines). Genes selected for RT-qPCR analysis are labeled. (C,D) RT-qPCR analysis comparing transcript levels of 6 DREAM target genes that were upregulated in RNA-seq (C) and 6 DREAM target genes that were not upregulated in RNA-seq (D). Transcript levels in late embryos are from wild type N2 (white), *lin-35* (black), *lin-52(WT)* (white), *lin-52(1A)* (light grey), and *lin-52(3A)* (dark grey). Expression values from 6 biological replicates were averaged and are presented as the relative quantity compared to *act-2*. Error bars indicate standard error of the mean, fold change values greater than 1.5-fold are provided, and significance was determined by a student’s t-test between transcript levels in mutant (3A or 1A) versus WT or *lin-35* versus N2 (* p-value < 0.05, ** p-value < 0.01).

To directly compare at one developmental stage the effect on transcription of loss of LIN-35 versus severing the LIN-35-MuvB association, we assessed transcript levels of a subset of DREAM target genes in *N2, lin-35(n745), lin-52(WT), lin-52(1A)*, and *lin-52(3A)* late embryos using RT-qPCR analysis (Figure 5C,D). We tested 6 DREAM target genes that were upregulated (Figure 5C) and 6 DREAM target genes that were not upregulated in *lin-52(3A)* late embryos (Figure 5D), as identified from our RNA-seq analysis. For the 6 DREAM target genes that were upregulated in our RNA-seq analysis (*C09G5*.*7, set-21, polh-1, ZK795*.*2, rnf-1*, and *Y47G6A*.*3)*, we consistently observed 1) modest but significant upregulation in *lin-52(3A)* versus *lin-52(WT) and* 2) a greater increase in transcript levels in *lin-35* versus N2 than in *lin-52(3A)* versus *lin-52(WT)* (Figure 5C). Consistent with our phenotype analysis, we observed that *lin-52(1A)* had a minimal effect on DREAM target gene levels (Figure 5C). For the 6 DREAM target genes that were not upregulated in our RNA-seq analysis (*cdk-1, air-1, mis-12, rad-51, kbp-3*, and *polk-1*), we did not observe an increase in transcript levels in *lin-52(3A)* or *lin-52(1A)* compared to *lin-52(WT)* (Figure 5D). Most (4 of 6) of these genes were significantly upregulated in *lin-35* versus N2 late embryos. Altogether, our RNA-seq and RT-qPCR analyses demonstrate that blocking DREAM assembly has a limited impact on DREAM target gene repression in late embryos, in terms of both number of genes upregulated and the amplitude of upregulation, as compared to complete loss of LIN-35.

### E2F-DP-LIN-35 and MuvB subcomplexes independently co-occupy chromatin sites

In the absence of LIN-35, E2F-DP and MuvB do not associate with one another and their chromatin occupancy is reduced genome-wide (Goetsch *et al*. 2017). In our *lin-52(3A)* worm strain, LIN-35 is present, but its association with MuvB is severed. We tested the impact of this severing on the chromatin localization of DREAM components using chromatin immunoprecipitation (ChIP). We chose 2 DREAM target genes that in our RNA-seq analysis were upregulated in *lin-52(3A)* (*set-21* and *polh-1*; Figure 6A), and 2 DREAM target genes that were not upregulated in *lin-52(3A)* (*air-1* and *mis-12*; Figure 6B). We performed ChIP-qPCR of DPL-1 and LIN-37 in *lin-35* null compared to N2 embryos, and DPL-1, LIN-35, LIN-37, and LIN-52 via its GFP tag in *lin-52(3A)* compared to *lin-52(WT)* embryos. IgG was used as a negative control for both ChIP analyses. We observed that all tested DREAM components remained similarly enriched at the 4 selected promoters in *lin-52(3A)* as compared to *lin-52(WT)* (Figure 6A,B). In contrast, chromatin occupancy of both DPL-1 and LIN-37 were significantly reduced in 3 of the 4 selected promoters in *lin-35(n745)* as compared to N2. An additional 8 DREAM target gene promoters were tested and showed similar DREAM chromatin occupancy profiles (Figure S3A,B). We did observe that LIN-37 and LIN-52 chromatin occupancy was reduced at some DREAM target gene promoters, but the observed decrease did not correspond to whether the target gene is misregulated (Figure S3A,B). Notably, the key difference between *lin-35(n745)* and *lin-52(3A)* is the status of the pocket protein.

**Figure 6.**
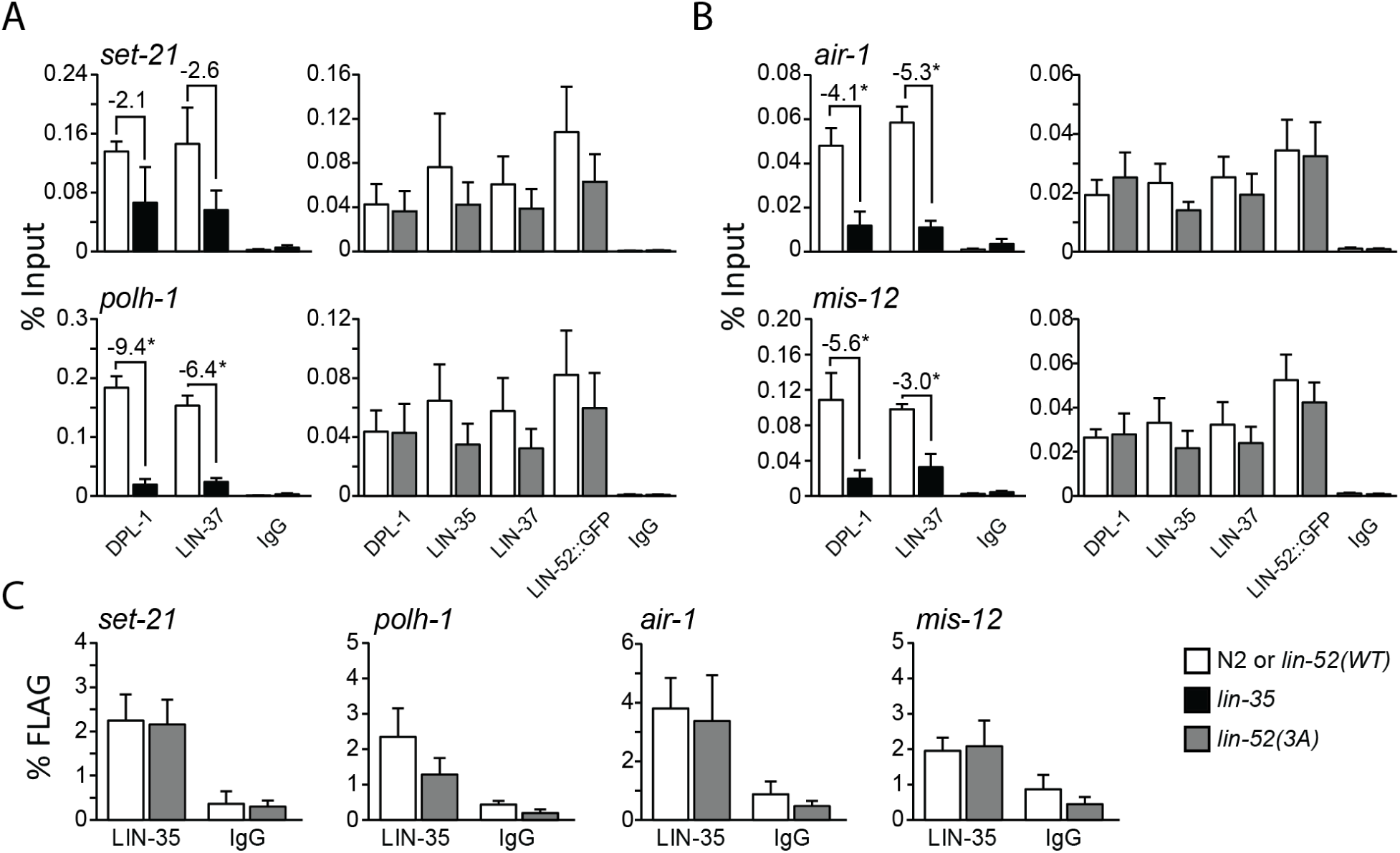
Analysis of chromatin localization of DREAM subunits at target genes. (A,B) ChIP-qPCR of DREAM subunits DPL-1 and LIN-37 in N2 (white) versus *lin-35* (black) and DPL-1, LIN-35, LIN-37, and LIN-52 (via GFP tag) in *lin-52(WT*) (white) versus *lin-52(3A*) (dark grey) late embryo extracts at 2 DREAM target genes that were upregulated in RNA-seq (A) and 2 DREAM target genes that were not upregulated in RNA-seq (B). IgG was used as a negative control. Signals are presented as percentage of Input DNA, with negative fold-change values greater than 2-fold noted. Error bars indicate standard error of the mean. Significance was determined by a student’s t-test between subunit ChIP values in mutant (*lin-35* or *lin-52*(*3A))* versus wild-type control (N2 or *lin-52(WT))* (* p-value < 0.05). (C) Sequential ChIP-qPCR of LIN-52 (via FLAG tag) followed by LIN-35 or IgG from *lin-52(WT*) (white) and *lin-52(3A*) (dark grey) late embryo extracts at 4 DREAM target genes. Signals are presented as percentage of FLAG IP DNA. Error bars indicate standard error of the mean. Additional data are shown in Figure S3.

Since DPL-1 and LIN-35 occupancy is not compromised by severing the LIN-52-LIN-35 association in *lin-52(3A)* while DPL-1 occupancy is compromised by loss of LIN-35, our results suggest that the chromatin association of the repressive E2F-DP transcription factor heterodimer is stabilized by its interaction with LIN-35.

We tested whether MuvB and E2F-DP-LIN-35 co-occupy DREAM target regions by performing sequential ChIP analysis. We first ChIPed LIN-52 via its FLAG tag and then ChIPed LIN-35. We observed no significant difference in LIN-52 and LIN-35 co-occupancy in *lin-52(3A)* extracts versus *lin-52(WT)* extracts at 8 DREAM target gene promoters, as determined by a student’s t-test (Figure 6C, S3C). Altogether, our results indicate that, although the interaction of LIN-35 and MuvB is disrupted, DREAM components can nevertheless co-localize at target promoters through their respective protein-DNA interactions.

## Discussion

We previously postulated that DREAM assembly, initiated by DYRK1A phosphorylation of LIN52 in mammalian cells (Litovchick *et al*. 2011; Guiley *et al*. 2015), stabilizes MuvB-mediated repression of DREAM target genes (Goetsch *et al*. 2017). Using CRISPR/Cas9-mediated targeted mutagenesis, we generated a mutant *C. elegans* strain in which LIN-52 was rendered incapable of interacting with LIN-35, the sole *C. elegans* Rb-like pocket protein. We expected that, similar to loss of LIN-35 (Goetsch *et al*. 2017), MuvB chromatin occupancy would be destabilized and its repressive function would be impaired. Instead, we observed that chromatin occupancy was not destabilized at the genes tested and that relatively few genes were upregulated. We observed that ∼2% of all DREAM target genes were de-repressed, indicating that for those genes MuvB chromatin occupancy alone is not sufficient for target gene repression. However, interrupting LIN-35-MuvB association did cause classic DREAM mutant phenotypes, including SynMuv, high temperature arrest, and ectopic expression of germline genes in the soma, consistent with impaired DREAM function. Altogether, we conclude that the LIN-35-MuvB association is not required to assemble DREAM at target sites but is necessary for full activity of the complex.

Our findings highlight that even minimal perturbation of DREAM-mediated transcriptional repression of target genes causes observable phenotypic consequences in *C. elegans*. Complete knockout of DREAM activity causes sterility, as demonstrated by our newly generated *lin-52* null mutant and null mutants of other DREAM components (Harrison *et al*. 2006). In contrast, although our RNA-seq analysis revealed that ∼2% of DREAM target genes were upregulated in the *lin-52(3A)* mutant, that mutant caused highly penetrant SynMuv phenotypes with no corresponding effect on fertility. Thus, phenotype analysis provides a sensitive readout of DREAM function.

How the limited upregulation of some DREAM target genes in *lin-52(3A)* mutant embryos causes the observed phenotypes remains unknown. The SynMuv phenotype is caused by ectopic activation of LIN-3/EGF in the hyp7 hypodermal syncytium, which requires synthetic impairment of redundant regulatory pathways established by SynMuv A and SynMuv B class genes (Cui *et al*. 2006). Although there is some evidence that SynMuv A class proteins directly target *lin-3*’s promoter (Saffer *et al*. 2011), DREAM does not appear to regulate *lin-3* directly (Goetsch *et al*. 2017). Similarly, we suspect that DREAM antagonism of high-temperature arrest and ectopic expression of germline genes in the soma is also indirect (Petrella *et al*. 2011; Goetsch *et al*. 2017; Rechtsteiner *et al*. 2019). Instead, we speculate that SynMuv B class proteins including DREAM regulate chromatin compaction in somatic cells during development (Costello and Petrella 2019). Thus, even minimal perturbation of DREAM complex activity may delay embryonic chromatin compaction, providing a window for alternative regulatory pathways to trigger the developmental defects detected at later larval stages.

Our analysis provides important insight into how assembly of the mammalian DREAM complex maintains repression of cell cycle genes and suggests that the pocket protein’s primary function is to protect MuvB’s function as a transcriptional repressor.

DREAM assembly is triggered by DYRK1A phosphorylation of LIN52, initiating MuvB association with p107/p130 (Litovchick *et al*. 2011; Guiley *et al*. 2015). Mammalian MuvB’s function switches from transcriptional repression in the DREAM complex during quiescence to transcriptional activation after associating with the B-Myb transcription factor and forming the Myb-MuvB (MMB) complex late in the cell cycle (Lewis *et al*. 2004; Osterloh *et al*. 2007; Schmit *et al*. 2007; Knight *et al*. 2009; Sadasivam *et al*. 2012). DYRK1A-mediated LIN52 phosphorylation also inhibits MuvB association with B-Myb (Litovchick *et al*. 2011), even though the 2 interaction interfaces do not physically overlap (Guiley *et al*. 2018). In *Drosophila*, Lin-52 protein is required for MMB to oppose repressive DREAM functions (Lewis *et al*. 2012), suggesting that Lin-52’s interaction with the pocket protein or B-Myb dictates MuvB’s transcriptional function (Guiley *et al*. 2018). Our data demonstrate that in worms MuvB localizes to chromatin sites and represses gene targets without direct association with the pocket protein.

These findings suggest that the pocket protein’s association with LIN-52 serves a different purpose than just to direct assembly of DREAM. We propose that the pocket protein’s association with LIN-52 instead primarily functions to oppose B-Myb association with MuvB, preventing MuvB from converting to its transcriptionally active role. Since no B-Myb homolog exists in *C. elegans* (Vorster *et al*. 2020), loss of direct LIN-35-MuvB association had little effect on target gene repression. We predict that a similar 3A mutation in LIN-52 in *Drosophila* or mammalian cells would cause significantly more upregulation of DREAM target genes.

The switch between MuvB-associated cell cycle gene repression and activation is hijacked in cancer cells. All 3 mammalian pocket proteins are inactivated by the E7 viral oncoprotein present in high-risk human papillovirus (HPV) (Zhang *et al*. 2006; Huh *et al*. 2007). E7 interacts with the mammalian pocket proteins through its high-affinity LxCxE binding motif, disrupting MuvB association with the pocket protein in DREAM (Guiley *et al*. 2015). HPV E7 concurrently coaxes MuvB into its transcriptional activator function by stimulating MMB assembly (Pang *et al*. 2014). However, cancer cells resist cytotoxic chemotherapy by temporarily exiting the cell cycle (Boichuk *et al*. 2013), suggesting that MuvB’s capacity for transcriptional repression is retained. Based on our findings that MuvB does not require direct association with the pocket protein to repress target genes, MuvB’s function in cancer cells requires closer scrutiny.

We previously observed that E2F-DP and MuvB chromatin association is severely affected by loss of LIN-35 (Goetsch *et al*. 2017). By severing LIN-35-MuvB association, the present study suggests a new model for DREAM complex formation where LIN-35 directly stabilizes E2F-DP chromatin occupancy. We also observed that MuvB chromatin occupancy is not disrupted even though MuvB no longer associates directly with E2F-DP-LIN-35. Importantly, *in vitro* analysis of the DNA-binding characteristics of heterodimeric mammalian E2F-DP identified a distinct induction of DNA bending, especially in the case of the homologues of *C. elegans* EFL-1-DPL-1 (E2F4-DP1/2) (Tao *et al*. 1997). We propose that DNA-associated E2F-DP heterodimers promote MuvB co-occupancy through a DNA bending-dependent mechanism. Together, our results suggest a model in which the LIN-35 pocket protein promotes E2F-DP chromatin occupancy, which in turn promotes MuvB chromatin occupancy.

Our results support an exciting model for how local E2F-DP-mediated alterations to DNA shape enhanced by their interaction with a pocket protein promote MuvB co-occupancy. Even with evolutionary divergence from the ancestral pocket protein, this model may also apply to pRb function. Many histone deacetylases and chromatin remodeling complexes associate with pRb through the LxCxE binding cleft, although many of these associations have only limited support thus far from structural/biochemical interaction studies (Dyson 2016). Variation in pRb monophosphorylation events that can alter pRb structure and recognition of binding partners offers one explanation for how pRb can potentially interact with >300 individual protein partners (Rubin 2013; Narasimha *et al*. 2014). Our data provide an alternative, but not exclusive, possibility, namely that direct and stable pRb association with these myriad protein partners may be unnecessary. Perhaps pRb association with a few partners such as E2F-DPs promotes localization of multi-protein complexes to genomic sites. Additional dissection of DREAM and pRb structure and function will shed light on how the pocket proteins mediate their essential cellular roles.

## Acknowledgments

We thank Seth Rubin and members of the Rubin and Strome labs for helpful discussions. Some strains were provided by the Caenorhabditis Genetics Center, which is funded by the NIH Office of Research Infrastructure Programs (P40 OD010440). This work was supported by National Institutes of Health R01 grant GM34069 to S.S., American Cancer Society Postdoctoral Fellowship PF-16-106-01-DDC to P.D.G., and National Institutes of Health R15 grant GM137145 to P.D.G.

## References

Anders, S., P. T. Pyl and W. Huber, 2015 HTSeq--a Python framework to work with high-throughput sequencing data. Bioinformatics 31: 166–169.

Arribere, J. A., R. T. Bell, B. X. Fu, K. L. Artiles, P. S. Hartman et al., 2014 Efficient marker-free recovery of custom genetic modifications with CRISPR/Cas9 in Caenorhabditis elegans. Genetics 198: 837–846.

Asthana, A., P. Ramanan, A. Hirschi, K. Z. Guiley, T. U. Wijeratne et al., 2022 The MuvB complex binds and stabilizes nucleosomes downstream of the transcription start site of cell-cycle dependent genes. Nat Commun 13: 526.

Beitel, G. J., E. J. Lambie and H. R. Horvitz, 2000 The C. elegans gene lin-9,which acts in an Rb-related pathway, is required for gonadal sheath cell development and encodes a novel protein. Gene 254: 253–263.

Boichuk, S., J. A. Parry, K. R. Makielski, L. Litovchick, J. L. Baron et al., 2013 The DREAM complex mediates GIST cell quiescence and is a novel therapeutic target to enhance imatinib-induced apoptosis. Cancer Res 73: 5120–5129.

Bolstad, B. M., R. A. Irizarry, M. Astrand and T. P. Speed, 2003 A comparison of normalization methods for high density oligonucleotide array data based on variance and bias. Bioinformatics 19: 185–193.

Boxem, M., and S. van den Heuvel, 2002 C. elegans class B synthetic multivulva genes act in G(1) regulation. Curr Biol 12: 906–911.

Burke, J. R., A. J. Deshong, J. G. Pelton and S. M. Rubin, 2010 Phosphorylation-induced conformational changes in the retinoblastoma protein inhibit E2F transactivation domain binding. J Biol Chem 285: 16286–16293.

Burkhart, D. L., and J. Sage, 2008 Cellular mechanisms of tumour suppression by the retinoblastoma gene. Nat Rev Cancer 8: 671–682.

Cao, L., B. Peng, L. Yao, X. Zhang, K. Sun et al., 2010 The ancient function of RB-E2F pathway: insights from its evolutionary history. Biol Direct 5: 55.

Ceol, C. J., F. Stegmeier, M. M. Harrison and H. R. Horvitz, 2006 Identification and classification of genes that act antagonistically to let-60 Ras signaling in Caenorhabditis elegans vulval development. Genetics 173: 709–726.

Chen, S., Y. Zhou, Y. Chen and J. Gu, 2018 fastp: an ultra-fast all-in-one FASTQ preprocessor. Bioinformatics 34: i884–i890.

Chi, W., and V. Reinke, 2006 Promotion of oogenesis and embryogenesis in the C. elegans gonad by EFL-1/DPL-1 (E2F) does not require LIN-35 (pRB). Development 133: 3147–3157.

Classon, M., and N. Dyson, 2001 p107 and p130: versatile proteins with interesting pockets. Exp Cell Res 264: 135–147.

Classon, M., and E. Harlow, 2002 The retinoblastoma tumour suppressor in development and cancer. Nat Rev Cancer 2: 910–917.

Cobrinik, D., 2005 Pocket proteins and cell cycle control. Oncogene 24: 2796–2809.

Costello, M. E., and L. N. Petrella, 2019 C. elegans synMuv B proteins regulate spatial and temporal chromatin compaction during development. Development 146.

Cui, M., J. Chen, T. R. Myers, B. J. Hwang, P. W. Sternberg et al., 2006 SynMuv genes redundantly inhibit lin-3/EGF expression to prevent inappropriate vulval induction in C. elegans. Dev Cell 10: 667–672.

Dick, F. A., and S. M. Rubin, 2013 Molecular mechanisms underlying RB protein function. Nat Rev Mol Cell Biol 14: 297–306.

Dickinson, D. J., A. M. Pani, J. K. Heppert, C. D. Higgins and B. Goldstein, 2015 Streamlined Genome Engineering with a Self-Excising Drug Selection Cassette. Genetics 200: 1035–1049.

Dickinson, D. J., J. D. Ward, D. J. Reiner and B. Goldstein, 2013 Engineering the Caenorhabditis elegans genome using Cas9-triggered homologous recombination. Nat Methods 10: 1028–1034.

Dobin, A., C. A. Davis, F. Schlesinger, J. Drenkow, C. Zaleski et al., 2013 STAR: ultrafast universal RNA-seq aligner. Bioinformatics 29: 15–21.

Dyson, N. J., 2016 RB1: a prototype tumor suppressor and an enigma. Genes Dev 30: 1492–1502.

Edelstein, A. D., M. A. Tsuchida, N. Amodaj, H. Pinkard, R. D. Vale et al., 2014 Advanced methods of microscope control using muManager software. J Biol Methods 1.

Fay, D. S., and J. Yochem, 2007 The SynMuv genes of Caenorhabditis elegans in vulval development and beyond. Dev Biol 306: 1–9.

Friedland, A. E., Y. B. Tzur, K. M. Esvelt, M. P. Colaiacovo, G. M. Church et al., 2013 Heritable genome editing in C. elegans via a CRISPR-Cas9 system. Nat Methods 10: 741–743.

Frokjaer-Jensen, C., M. W. Davis, C. E. Hopkins, B. J. Newman, J. M. Thummel et al., 2008 Single-copy insertion of transgenes in Caenorhabditis elegans. Nat Genet 40: 1375–1383.

Gibson, D. G., L. Young, R. Y. Chuang, J. C. Venter, C. A. Hutchison, 3rd et al., 2009 Enzymatic assembly of DNA molecules up to several hundred kilobases. Nat Methods 6: 343–345.

Goetsch, P. D., J. M. Garrigues and S. Strome, 2017 Loss of the Caenorhabditis elegans pocket protein LIN-35 reveals MuvB’s innate function as the repressor of DREAM target genes. PLoS Genet 13: e1007088.

Guiley, K. Z., A. N. Iness, S. Saini, S. Tripathi, J. S. Lipsick et al., 2018 Structural mechanism of Myb-MuvB assembly. Proc Natl Acad Sci U S A 115: 10016–10021.

Guiley, K. Z., T. J. Liban, J. G. Felthousen, P. Ramanan, L. Litovchick et al., 2015 Structural mechanisms of DREAM complex assembly and regulation. Genes Dev 29: 961–974.

Harrison, M. M., C. J. Ceol, X. Lu and H. R. Horvitz, 2006 Some C. elegans class B synthetic multivulva proteins encode a conserved LIN-35 Rb-containing complex distinct from a NuRD-like complex. Proc Natl Acad Sci U S A 103: 16782–16787.

Harrison, M. M., X. Lu and H. R. Horvitz, 2007 LIN-61, one of two Caenorhabditis elegans malignant-brain-tumor-repeat-containing proteins, acts with the DRM and NuRD-like protein complexes in vulval development but not in certain other biological processes. Genetics 176: 255–271.

Helin, K., J. A. Lees, M. Vidal, N. Dyson, E. Harlow et al., 1992 A cDNA encoding a pRB-binding protein with properties of the transcription factor E2F. Cell 70: 337–350.

Huang, L. S., P. Tzou and P. W. Sternberg, 1994 The lin-15 locus encodes two negative regulators of Caenorhabditis elegans vulval development. Mol Biol Cell 5: 395–411.

Huh, K., X. Zhou, H. Hayakawa, J. Y. Cho, T. A. Libermann et al., 2007 Human papillomavirus type 16 E7 oncoprotein associates with the cullin 2 ubiquitin ligase complex, which contributes to degradation of the retinoblastoma tumor suppressor. J Virol 81: 9737–9747.

Hurford, R. K., Jr., D. Cobrinik, M. H. Lee and N. Dyson, 1997 pRB and p107/p130 are required for the regulated expression of different sets of E2F responsive genes. Genes Dev 11: 1447–1463.

Irizarry, R. A., B. Hobbs, F. Collin, Y. D. Beazer-Barclay, K. J. Antonellis et al., 2003 Exploration, normalization, and summaries of high density oligonucleotide array probe level data. Biostatistics 4: 249–264.

Kim, E., L. Sun, C. V. Gabel and C. Fang-Yen, 2013 Long-term imaging of Caenorhabditis elegans using nanoparticle-mediated immobilization. PLoS One 8: e53419.

Kirienko, N. V., and D. S. Fay, 2007 Transcriptome profiling of the C. elegans Rb ortholog reveals diverse developmental roles. Dev Biol 305: 674–684.

Knight, A. S., M. Notaridou and R. J. Watson, 2009 A Lin-9 complex is recruited by B-Myb to activate transcription of G2/M genes in undifferentiated embryonal carcinoma cells. Oncogene 28: 1737–1747.

Korenjak, M., B. Taylor-Harding, U. K. Binne, J. S. Satterlee, O. Stevaux et al., 2004 Native E2F/RBF complexes contain Myb-interacting proteins and repress transcription of developmentally controlled E2F target genes. Cell 119: 181–193.

Lees, J. A., M. Saito, M. Vidal, M. Valentine, T. Look et al., 1993 The retinoblastoma protein binds to a family of E2F transcription factors. Mol Cell Biol 13: 7813–7825.

Lewis, P. W., E. L. Beall, T. C. Fleischer, D. Georlette, A. J. Link et al., 2004 Identification of a Drosophila Myb-E2F2/RBF transcriptional repressor complex. Genes Dev 18: 2929–2940.

Lewis, P. W., D. Sahoo, C. Geng, M. Bell, J. S. Lipsick et al., 2012 Drosophila lin-52 acts in opposition to repressive components of the Myb-MuvB/dREAM complex. Mol Cell Biol 32: 3218–3227.

Liban, T. J., E. M. Medina, S. Tripathi, S. Sengupta, R. W. Henry et al., 2017 Conservation and divergence of C-terminal domain structure in the retinoblastoma protein family. Proc Natl Acad Sci U S A 114: 4942–4947.

Liban, T. J., M. J. Thwaites, F. A. Dick and S. M. Rubin, 2016 Structural Conservation and E2F Binding Specificity within the Retinoblastoma Pocket Protein Family. J Mol Biol 428: 3960–3971.

Litovchick, L., L. A. Florens, S. K. Swanson, M. P. Washburn and J. A. DeCaprio, 2011 DYRK1A protein kinase promotes quiescence and senescence through DREAM complex assembly. Genes Dev 25: 801–813.

Litovchick, L., S. Sadasivam, L. Florens, X. Zhu, S. K. Swanson et al., 2007 Evolutionarily conserved multisubunit RBL2/p130 and E2F4 protein complex represses human cell cycle-dependent genes in quiescence. Mol Cell 26: 539–551.

Love, M. I., W. Huber and S. Anders, 2014 Moderated estimation of fold change and dispersion for RNA-seq data with DESeq2. Genome Biol 15: 550.

Lu, X., and H. R. Horvitz, 1998 lin-35 and lin-53, two genes that antagonize a C. elegans Ras pathway, encode proteins similar to Rb and its binding protein RbAp48. Cell 95: 981–991.

Mages, C. F., A. Wintsche, S. H. Bernhart and G. A. Muller, 2017 The DREAM complex through its subunit Lin37 cooperates with Rb to initiate quiescence. Elife 6.

Marceau, A. H., J. G. Felthousen, P. D. Goetsch, A. N. Iness, H. W. Lee et al., 2016 Structural basis for LIN54 recognition of CHR elements in cell cycle-regulated promoters. Nat Commun 7: 12301.

Marnik, E. A., J. H. Fuqua, C. S. Sharp, J. D. Rochester, E. L. Xu et al., 2019 Germline Maintenance Through the Multifaceted Activities of GLH/Vasa in Caenorhabditis elegans P Granules. Genetics 213: 923–939.

Muller, G. A., and K. Engeland, 2010 The central role of CDE/CHR promoter elements in the regulation of cell cycle-dependent gene transcription. FEBS J 277: 877–893.

Muller, G. A., M. Quaas, M. Schumann, E. Krause, M. Padi et al., 2012 The CHR promoter element controls cell cycle-dependent gene transcription and binds the DREAM and MMB complexes. Nucleic Acids Res 40: 1561–1578.

Muller, G. A., K. Stangner, T. Schmitt, A. Wintsche and K. Engeland, 2017 Timing of transcription during the cell cycle: Protein complexes binding to E2F, E2F/CLE, CDE/CHR, or CHR promoter elements define early and late cell cycle gene expression. Oncotarget 8: 97736–97748.

Myers, T. R., and I. Greenwald, 2005 lin-35 Rb acts in the major hypodermis to oppose ras-mediated vulval induction in C. elegans. Dev Cell 8: 117–123.

Narasimha, A. M., M. Kaulich, G. S. Shapiro, Y. J. Choi, P. Sicinski et al., 2014 Cyclin D activates the Rb tumor suppressor by mono-phosphorylation. Elife 3.

Osterloh, L., B. von Eyss, F. Schmit, L. Rein, D. Hubner et al., 2007 The human synMuv-like protein LIN-9 is required for transcription of G2/M genes and for entry into mitosis. EMBO J 26: 144–157.

Pang, C. L., S. Y. Toh, P. He, S. Teissier, Y. Ben Khalifa et al., 2014 A functional interaction of E7 with B-Myb-MuvB complex promotes acute cooperative transcriptional activation of both S-and M-phase genes. (129 c). Oncogene 33: 4039–4049.

Petrella, L. N., W. Wang, C. A. Spike, A. Rechtsteiner, V. Reinke et al., 2011 synMuv B proteins antagonize germline fate in the intestine and ensure C. elegans survival. Development 138: 1069–1079.

Pilkinton, M., R. Sandoval and O. R. Colamonici, 2007 Mammalian Mip/LIN-9 interacts with either the p107, p130/E2F4 repressor complex or B-Myb in a cell cycle-phase-dependent context distinct from the Drosophila dREAM complex. Oncogene 26: 7535–7543.

Rechtsteiner, A., M. E. Costello, T. A. Egelhofer, J. M. Garrigues, S. Strome et al., 2019 Repression of Germline Genes in Caenorhabditis elegans Somatic Tissues by H3K9 Dimethylation of Their Promoters. Genetics 212: 125–140.

Rubin, S. M., 2013 Deciphering the retinoblastoma protein phosphorylation code. Trends Biochem Sci 38: 12–19.

Sadasivam, S., S. Duan and J. A. DeCaprio, 2012 The MuvB complex sequentially recruits B-Myb and FoxM1 to promote mitotic gene expression. Genes Dev 26: 474–489.

Saffer, A. M., D. H. Kim, A. van Oudenaarden and H. R. Horvitz, 2011 The Caenorhabditis elegans synthetic multivulva genes prevent ras pathway activation by tightly repressing global ectopic expression of lin-3 EGF. PLoS Genet 7: e1002418.

Schmit, F., S. Cremer and S. Gaubatz, 2009 LIN54 is an essential core subunit of the DREAM/LINC complex that binds to the cdc2 promoter in a sequence-specific manner. FEBS J 276: 5703–5716.

Schmit, F., M. Korenjak, M. Mannefeld, K. Schmitt, C. Franke et al., 2007 LINC, a human complex that is related to pRB-containing complexes in invertebrates regulates the expression of G2/M genes. Cell Cycle 6: 1903–1913.

Schneider, C. A., W. S. Rasband and K. W. Eliceiri, 2012 NIH Image to ImageJ: 25 years of image analysis. Nat Methods 9: 671–675.

Tabuchi, T. M., B. Deplancke, N. Osato, L. J. Zhu, M. I. Barrasa et al., 2011 Chromosome-biased binding and gene regulation by the Caenorhabditis elegans DRM complex. PLoS Genet 7: e1002074.

Tao, Y., R. F. Kassatly, W. D. Cress and J. M. Horowitz, 1997 Subunit composition determines E2F DNA-binding site specificity. Mol Cell Biol 17: 6994–7007.

Tedesco, D., J. Lukas and S. I. Reed, 2002 The pRb-related protein p130 is regulated by phosphorylation-dependent proteolysis via the protein-ubiquitin ligase SCF(Skp2). Genes Dev 16: 2946–2957.

Vorster, P. J., P. Goetsch, T. U. Wijeratne, K. Z. Guiley, L. Andrejka et al., 2020 A long lost key opens an ancient lock: Drosophila Myb causes a synthetic multivulval phenotype in nematodes. Biol Open 9.

Wang, D., S. Kennedy, D. Conte, Jr., J. K. Kim, H. W. Gabel et al., 2005 Somatic misexpression of germline P granules and enhanced RNA interference in retinoblastoma pathway mutants. Nature 436: 593–597.

Zhang, B., W. Chen and A. Roman, 2006 The E7 proteins of low-and high-risk human papillomaviruses share the ability to target the pRB family member p130 for degradation. Proc Natl Acad Sci U S A 103: 437–442.

Zwicker, J., F. C. Lucibello, L. A. Wolfraim, C. Gross, M. Truss et al., 1995 Cell cycle regulation of the cyclin A, cdc25C and cdc2 genes is based on a common mechanism of transcriptional repression. EMBO J 14: 4514–4522.

