## Supplemental Information for "DREAM Interrupted: Severing LIN-35-MuvB association in *Caenorhabditis elegans* impairs DREAM function but not its chromatin localization"

**Supplemental Materials and Methods:**

**CRISPR/Cas9-mediated genome editing**

To generate *lin-52(KO),* 2 Cas9 target sites were identified near the 5' and 3' ends of the *lin-52 gene*. Single guide RNA sequences were cloned into the PU6::unc119_sgRNA vector. The *lin-52* KO homologous repair template was generated by amplifying homology arms containing the *lin-52* promoter and *lin-52* 3' UTR and cloned into the N-terminal tag digested pDD284 vector. The following injection mix was microinjected into the germline of ~50 N2 young adults: 50 ng / µL Cas9 expression plasmid (pDD162, Addgene #47549), 2.5 ng / µL *Pmyo-2::mCherry::unc-54utr* (pCJF90, Addgene #19327), 5 ng / µL *Pmyo-3::mCherry::unc-54utr* (pCFJ104, Addgene #19328), 10 ng / µL *Prab-3::mCherry::unc-54utr* (pGH8, Addgene #19359), 50 ng / µL *lin-52* 5' sgRNA (pPDG14), 50 ng / µL *lin-52* 3' sgRNA (pPDG18), and 10 ng µL *Plin-52::TagRFP-T^SEC^3xFLAG::lin-52utr* (pPDG13). CRISPR/Cas9-positive progeny were treated with hygromycin and screened for the Roller phenotype and absence of fluorescent co-injection marker expression (the latter enables extrachromosomal arrays to be distinguished from edited endogenous genes). Individuals from 1 positive selection plate were selected and balanced to create the strain SS1240 *lin-52(bn132(lin-52p::TagRFP-T^SEC^3xFLAG::lin-52 3' UTR)) III / hT2G [bli-4(e937) let-?(q782) qIs48] (I:III)*. The self-excising cassette (SEC) was removed by a 4-5 hour heat-shock of L1 larvae at 32°C. Non-Roller F1 progeny were isolated to create the strain SS1241 *lin-52(bn133(lin-52p::TagRFP-T::3xFLAG::lin-52 3' UTR)) III / hT2G [bli-4(e937) let-?(q782) qIs48] (I:III).*

To generate *lin-52(WT)*, 2 Cas9 target sites were identified near the 5' and 3' ends of the *TagRFP-T-3xFLAG* coding sequence. Single guide RNA sequences were cloned into pDD162. The *lin-52* WT homologous repair template was generated by amplifying homology arms containing the *lin-52* promoter with the gene's coding sequence and the *lin-52* 3' UTR and cloned into the C-terminal tag digested pDD282 vector. The following CRISPR/Cas9 and co-injection marker plasmid mix was microinjected into the germline of ~50 SS1241 young adults: 50 ng / µL *TagRFP-T* 5' sgRNA-Cas9 vector (pPDG21), 50 ng / µL *TagRFP-T* 3' sgRNA-Cas9 vector (pPDG22), 2.5 ng / µL pCJF90, 5 ng / µL pCFJ104, and 10 ng / µL *Plin-52::lin-52 CDS-GFP^SEC^3xFLAG::lin-52utr* (pPDG17). CRISPR/Cas9-positive progeny were treated with hygromycin and screened for the Roller phenotype and absence of fluorescent co-injection marker expression. Individuals from 2 of 3 positive selection plates were selected and made homozygous to create strains SS1325 and SS1326 *lin-52(bn138(lin-52::GFP^SEC^3xFLAG)) III*. The SEC was removed by heat-shock, and non-Roller F1 progeny were isolated to create the strains SS1256 and SS1257 *lin-52(bn139(lin-52::GFP::3xFLAG)) III*. SS1256 was backcrossed 6 times to generate strain SS1272, which was used in downstream experiments.

To generate *lin-52(1A)* and *lin-52(3A)*, 1 Cas9 target site was identified near the LxCxE coding sequence and cloned into the pDD162 vector. Single strand DNA templates included at least 40 base pairs of homology flanking the LxCxE coding sequence and silent mutations to aid in genotyping, as illustrated in Figure 2B. The following/Cas9 and co-injection marker plasmid mix was microinjected into the germline of 6 (for 1A) and 10 (for 3A) SS1256 young adults: 40 ng / µL *lin-52* LxCxE sgRNA-

Cas9 vector (pPDG59), 2.5 ng / µL pCJF90, 5 ng / µL pCFJ104, 20 ng / µL *lin-52* mutagenesis ssDNA template (1A or 3A), 40 ng / µL *dpy-10(cn64)* sgRNA (pJA58, Addgene plasmid #59933), and *dpy-10(cn64)* ssDNA template. *dpy-10(cn64)* guide and ssDNA template were co-injected to select for positive CRISPR activity in injectant progeny. Injected adults were cloned onto individual plates, and F1 progeny were screened for presence of a Roller (Rol) and/or Dumpy (Dpy) phenotype. Individual Rol and/or Dpy progeny were genotyped, resulting in 3 independent *lin-52(1A*) and 2 independent *lin-52(3A)* strains. Each strain was backcrossed 6 times to create SS1273-SS1275 *lin-52(bn150(lin-52[C44A]::GFP::3xFLAG)) III,* and SS1276 and SS1277 *lin-52(bn151(lin-52[L42A,C44A,E46A]::GFP::3xFLAG)) III*. SS1273 and SS1276 were used in downstream experiments.

**Immunoblotting and co-immunoprecipitation (coIP)**

CoIP lysates were prepared by grinding frozen embryos using a mortar and pestle, resuspending in lysis buffer (25 mM HEPES pH 7.6, 150 mM NaCl, 1mM DTT, 1mM EDTA, 0.5 mM EGTA, 0.1% Nonidet P-40, 10% glycerol) with Complete EDTA-free Protease Inhibitors (Roche), and sonicating twice for 30 seconds. Lysates were clarified and precleared using a mix of Protein A and Protein G Dynabeads (ThermoFisher). Protein concentrations of coIP lysates were determined using a Qubit fluorometer (ThermoFisher). For each IP, 5 µg of anti-FLAG was crosslinked to Protein G Dynabeads and 2 µg of anti-GFP or anti-LIN-35 was crosslinked to Protein A Dynabeads using dimethyl pimelimidate in 0.2 M trimethylamine pH 8.2. Crosslinking was stopped using 0.1M Tris pH 8.0, and beads were washed with 0.1 M glycine pH 2.8 before being stored in phosphate buffered saline pH 7.2 with 0.05% Tween-20. Each IP was washed with lysis buffer, and eluted with 50 µL 2x SDS gel-loading buffer for 5 minutes at 98°C

**Chromatin immunoprecipitation (ChIP) and sequential ChIP**

For ChIP, chromatin extracts were precleared with Protein A Dynabeads. ChIPs were performed with 2 mg of extract and 1 µg of antibody, with 2% of the extract set aside for an input reference control. ChIPs were incubated overnight at 4°C with 1% sarkosyl. Protein A Dynabeads equilibrated in 20 µL FA buffer were added and incubated for 2 hours at 4°C. ChIPs were washed with the following buffers: once with FA buffer containing 1 M NaCl, once with FA buffer containing 0.5 M NaCl, once with TEL buffer (10 mM Tris-HCl pH 8.0, 0.25 M LiCl, 1% NP-40, 1% sodium deoxycholate, 1 mM EDTA), and twice with TE buffer (10 mM Tris-HCl pH 8.0 and 1 mM EDTA). 2 elutions of 50 µL elution buffer containing TE plus 1% SDS and 250 mM NaCl were incubated at 55°C. Eluted ChIP and input samples were incubated with proteinase K for 1 hour at 55°C. Crosslinks were reversed overnight at 65°C. DNA was purified by phenol-chloroform extraction and ethanol precipitation using glycogen as a carrier.

For sequential ChIP, chromatin extracts were precleared with Protein G Dynabeads and 4 parallel ChIPs per replicate were performed with 2.5 mg of extract and 2.5 µg of anti-FLAG antibody, with 2% of the extract set aside for an input reference control. ChIPs were incubated overnight at 4°C with 1% sarkosyl. Protein G Dynabeads equilibrated in 20 µL FA buffer were added and incubated for 2 hours at 4°C. ChIPs for each replicate were washed as described above and pooled. 2 elutions of 50 µL 0.1M $NaHCO_3$ plus 1% SDS were incubated at 55°C for 15 minutes. Elutions were divided, diluted with FA buffer with 1% sarkosyl, and incubated with anti-LIN-35 or IgG as a negative control, with 10% of the elution set aside as a reference control. The 2nd ChIP was incubated overnight at 4°C. Protein A Dynabeads equilibrated in 20 μL FA buffer were added and incubated for 2 hours at 4°C. ChIPs were washed and eluted twice with 50 μL elution buffer with incubation at 55°C. Eluted ChIP, reference, and input samples were incubated with proteinase K for 1 hour at 55°C. Crosslinks were reversed overnight at 65°C. DNA was purified by phenol-chloroform extraction and ethanol precipitation using glycogen as a carrier.

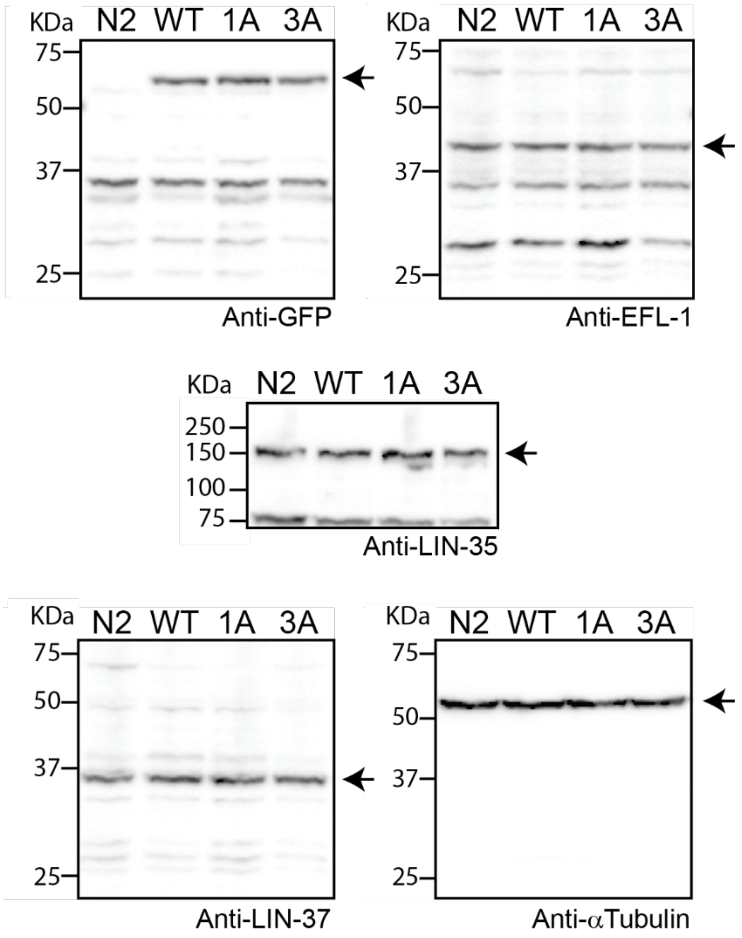

**Figure S1**

Full western blots of DREAM subunits LIN-52 (via GFP tag), EFL-1, LIN-35, and LIN-37 using whole worm lysates from Bristol (N2), *lin-52(WT), lin-52(1A),* and *lin-52(3A)* separated by SDS/PAGE. Antibodies used are indicated below each blot. Alpha-tubulin was used as a loading control. Membranes were cut at the 75 kDa band. Arrows indicate blot regions presented in Figure 2D.

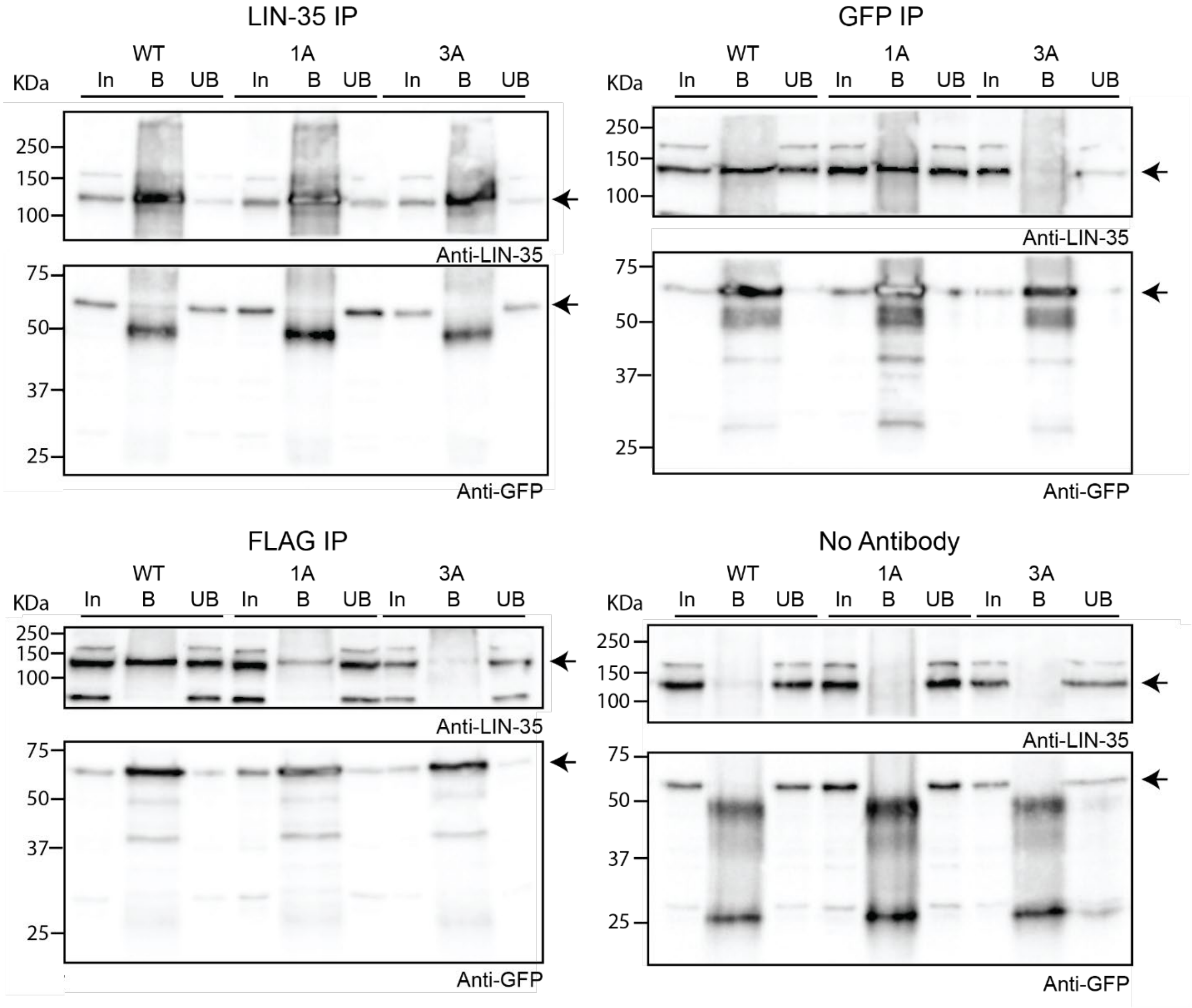

**Figure S2**

Full western blots of late embryo extracts from *lin-52(WT)*, *lin-52(1A)*, and *lin-52(3A)* that were immunoprecipitated with anti-LIN-35, anti-GFP, and anti-FLAG antibodies, with no antibody serving as a negative control. Proteins bound (B) and unbound (UB) were separated by SDS/PAGE and transferred to PVDF membranes that were cut at the 75 kDa band. Antibodies used are indicated below each blot. 5% of Input (In) was included. Arrows indicate blot regions presented in Figure 3.

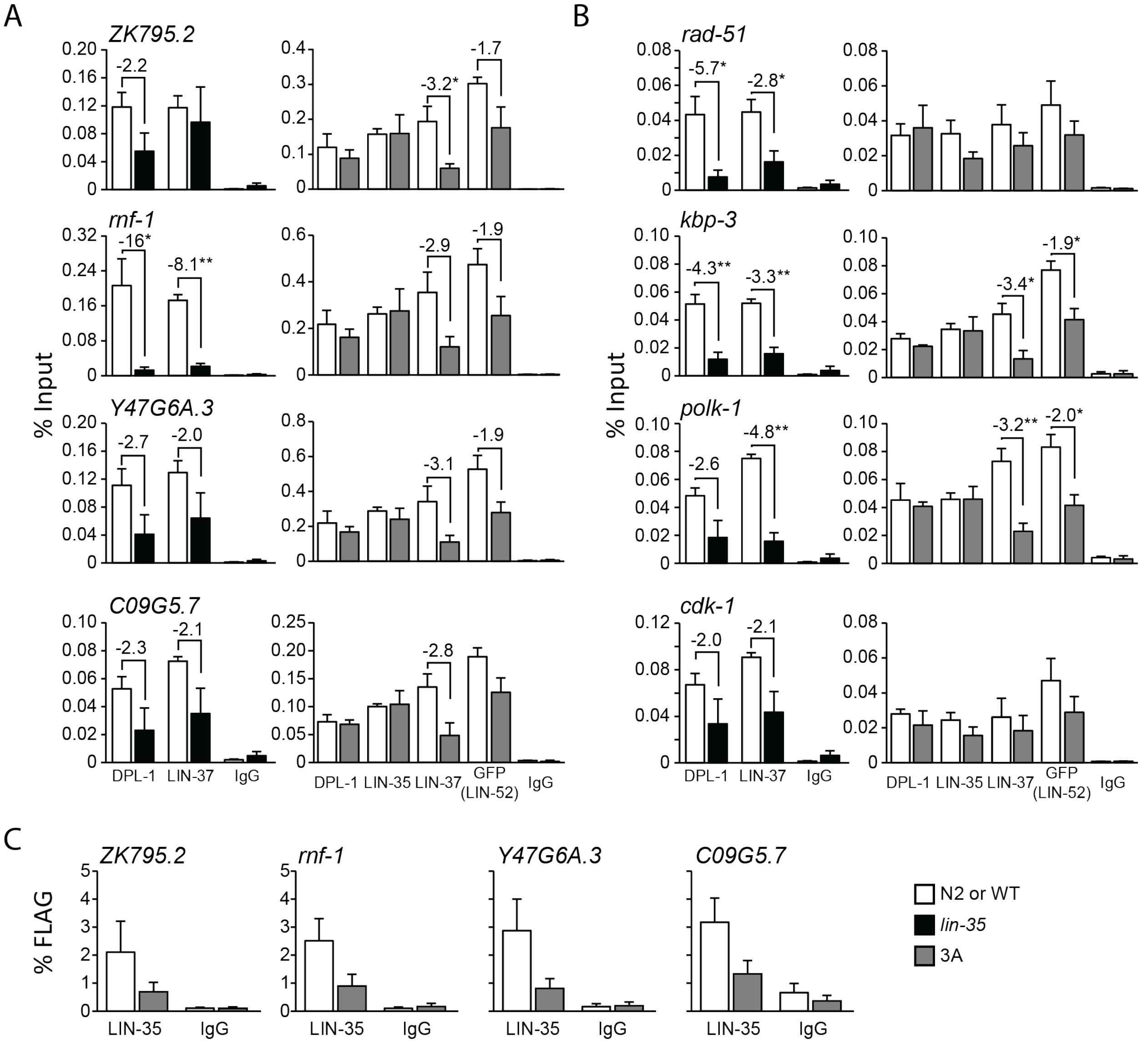

### Figure S3

(A) ChIP-qPCR of DREAM subunits DPL-1 and LIN-37 in N2 (white) versus *lin-35* (black) and DPL-1, LIN-35, LIN-37, and LIN-52 (via GFP tag) in *lin-52(WT)* (white) versus *lin-52(3A)* (dark grey) late embryo extracts at 8 DREAM target genes in addition to those shown in Figure 6: 4 that were upregulated in RNA-seq (A) and 4 that were not upregulated in RNA-seq (B). IgG was used as a negative control. Signals are presented as percentage of Input DNA, with negative fold-change values greater than 2-fold noted. Error bars indicate standard error of the mean. Significance was determined by a student's t-test between subunit ChIP values in mutant (*lin-35* or *lin-52(3A)*) versus wild-type control (N2 or *lin-52(WT)*) (* p-value < 0.05). (C) Sequential ChIP-qPCR of LIN-52 (via FLAG tag) followed by LIN-35 or IgG from *lin-52(WT)* (white) and *lin-52(3A)* (dark grey) late embryo extracts at 4 DREAM target genes in addition to those shown in Figure 6. Signals are presented as percentage of FLAG IP DNA. Error bars indicate standard error of the mean.

**Table S1**

| Antibody information | | | | |
|---|---|---|---|---|
| Protein | Antibody | Species | Type | Source |
| GFP | NB600-308 | Rabbit | Polyclonal | Novus |
| FLAG | M2 (F3165) | Mouse | Monoclonal | Sigma-Aldrich |
| DPL-1 | SDQ3599 | Rabbit | Polyclonal | SDIX/Novus |
| EFL-1 | SDQ3590 | Rabbit | Polyclonal | SDIX/Novus |
| LIN-35 (ChIP) | SDQ2003 | Rabbit | Polyclonal | SDIX/Novus |
| LIN-35 (CoIP/Western) | SDQ3232 | Rabbit | Polyclonal | SDIX/Novus |
| LIN-37 | SDQ3166 | Rabbit | Polyclonal | SDIX/Novus |
| Alpha-Tubulin | DM1A | Mouse | Monoclonal | - |
| Pre-immune serum (IgG) | - | - | - | - |

**Table S2**

| Name | Sequence | Notes |
|---|---|---|
| Cloning Primers | | |
| lin-52 gRNA 5' F | GTCGTATCCAATAAATCCTAGGTTTTAGAG CTAGAAATAGCAAGTTA | For pPDG14 |
| lin-52 gRNA 5' R | CTAGGATTTATTGGATACGACAAACATTTAG ATTTGCAATTCAATTATATAG | For pPDG14 |
| lin-52 gRNA 3' F | GAAGCCAGTGAATTGAATAGGTTTTAGAGC TAGAAATAGCAAGTTA | For pPDG18 |
| lin-52 gRNA 3' R | CTATTCAATTCACTGGCTTCAAACATTTAGA TTTGCAATTCAATTATATAG | For pPDG18 |
| lin-52_primer1_RFP | GTCACGACGTTGTAAAACGACGGCCAGTC GCATTCGAGCAAACCGGAGGA | For pPDG13 |
| lin-52_primer2_RFP_Nterm | CTTGATGAGCTCCTCTCCCTTGGAGACCAT TTTTTTCCTGAAATTACCGCTATATGTC | For pPDG13 |
| lin-52_primer3 | CGTGATTACAAGGATGACGATGACAAGAGA ATTGAATAGTGGTCTATCAAAAAATAATG | For pPDG13 and pPDG17 |
| lin-52_primer4_N-term | TCACACAGGAAACAGCTATGACCATGTTAT CACCTTGGGTACTTGCTGGAT | For pPDG13 |
| lin-52_primer1_GFP | ACGTTGTAAAACGACGGCCAGTCGCCGGC ACATTCGAGCAAACCGGAGGA | For pPDG17 |
| lin-52_primer2_C-term | CATCGATGCTCCTGAGGCTCCCGATGCTCC CTGGCTTCCTGTCGTTTCTTC | For pPDG17 |
| lin-52_primer4_C-term | GGAAACAGCTATGACCATGTTATCGATTTC CACCTTGGGTACTTGCTGGAT | For pPDG17 |
| sgRNA SDM R | CAAGACATCTCGCAATAGG | Reverse primer for Q5 targeted mutagenesis, see Dickinson et al, 2013 |
| tagRFP-sgRNA SDM 5' F | TGGCTTTCCCTCTCCCTCGGGTTTTAGAGC TAGAAATAGCAAGT | For pPDG21 |
| tagRFP-sgRNA SDM 3' F | TGTGTCCGAGCTTGGATGGGGTTTTAGAGC TAGAAATAGCAAGT | For pPDG22 |
| lin-52 LxCxE sgRNA SDM F | ACTTTCTTCTGGCATTGTTTGTTTTAGAGCT AGAAATAGCAAGT | For pPDG59 |
| lin-52 5’ check F | GAACAGGCGAAAATGCTTGGT | Genotyping and sequencing primer |
| lin-52 5’ check R | CAGTCACAGCATCTTCCTTGAGA | Genotyping and sequencing primer |
| ChIP-qPCR Primers | | |
| set-21 Pro F | ACGACGGGCCCAAAAGTAAA | |
| set-21 Pro R | TGTTGTTTCGTTTTCGCGAATTT | |
| polh-1 Pro F | TCAATGTTTGAAACCCCGCC | |
| polh-1 Pro R | ATACTCAGCCAAGCAGCCAA | |
| air-1 Pro F | ATTCGCAACGTGTCAGCAAC | |
| air-1 Pro R | ATGAATTTTGCTTGGCGGGT | |
| mis-12 Pro F | TTCCCGACAATTCGCTCTCC | |
| mis-12 Pro R | CGTGTATGCACACCTCACCT | |
| ZK795.2 Pro F | ATTCCATGAGTTTCAACGAAATACT | |
| ZK795.2 Pro R | TGTGCGTCAGATGAAATGCC | |
| rnf-1 Pro F | GCGTAGCATGTGACTTGTCG | |
| rnf-1 Pro R | GCGCTCCAATGCGAATTTCT | |

| | |
|---|---|
| Y47G6A.3 Pro F | TCCAATTGGCGGGGATTCAA |
| Y47G6A.3 Pro R | TGCACAAAACCTGTGTGAAAGT |
| C09G5.7 Pro F | TGTCGTGAAAAATGCGCTGG |
| C09G5.7 Pro R | TGGTCGAGTTTTCCCCTCTC |
| rad-51 Pro F | GCGCACTTGCTGTACTCTTG |
| rad-51 Pro R | CCGTTCCTATCGGTGCCTTT |
| kbp-3 Pro F | GTCAAGAGGGCCAAGGTCTC |
| kbp-3 Pro R | AATCGACTCAGTGCGAGAGG |
| polk-1 Pro F | CGCGGAGAAACTCCACTGAT |
| polk-1 Pro R | TTCTCGCGCACTCAACTTCT |
| cdk-1 Pro F | ACAATCCTTCTCAGCGCGT |
| cdk-1 Pro R | CGATAGAAAAGGCGTAAGCGTG |
| RT-qPCR primers | |
| act-2 RTref F | CGTCATCAAGGAGTCATGGTC |
| act-2 RTref F | CATGTCGTCCCAGTTGGTAA |
| pgl-1 1155-1238 F | ACGAGCCATTGCGGAACTTA |
| pgl-1 1155-1238 R | TGAGGAATCAGTGGAACGGG |
| pgl-3 1613-1700 F | CCATCAACTGCCTCACCTGT |
| pgl-3 1613-1700 R | AAGTCGAGATCTTCGGTGGC |
| glh-1 2415-2589 F | TCGGAAGAACTGGAAGAGTTGG |
| glh-1 2415-2589 R | ACACTGGACCCAAATCCACT |
| C09G5.7 974-1122 F | GCAAAAGTGTCCCACCATCG |
| C09G5.7 974-1122 R | TGAGTGAACTGGACGACGAC |
| set-21 1359-1524 F | AAATGTTGCGCGAACTGTCG |
| set-21 1359-1524 R | GTCCGTGTACGTCTTTCCGT |
| polh-1 759-842 F | TGTTCGAGGATTTGGCGGAA |
| polh-1 759-842 R | TCCACTTCGAGCAGTTCACC |
| ZK795.2 300-549 F | GTGTGATCCTGAGTGGGGAC |
| ZK795.2 300-549 R | TTTTTCTTCGCGTTCGGCTG |
| rnf-1 939-1121 F | ACTTCCGGCAGTTTTGGGAA |
| rnf-1 939-1121 R | GGAAAAGTCGAGGTTCCCGT |
| Y47G6A.3 368-567 F | AGTTGCTCAAAGAGCCCAGT |
| Y47G6A.3 368-567 R | CGCAGAAGACTTTGGCGTTC |
| cdk-1 703-911 F | TTCAGAGTTCTCGGCACACC |
| cdk-1 703-911 R | TTCGCGTTGAGACGAAGTGA |
| air-1 594-740 F | ACGCCATACATTGTGCGGTA |
| air-1 594-740 R | CCAGTTTGATTGGCGAACGG |
| mis-12 370-515 F | ATTCGACAGCTCCGCATCAA |
| mis-12 370-515 R | ATTCGTGTTGGGCTATCGGG |
| rad-51 (534-686) F | CAATGCCACTTTTCGACCCG |
| rad-51 (534-686) R | TCGGACATCATTGCTCCTGC |
| kbp-3 158-346 F | GCCACGAGCAACAGTCATTC |
| kbp-3 158-346 R | TCGGCGTGGTTTTCAAGAGA |
| polk-1 741-975 F | GTTCGGAACTGGACGAGAGG |
| polk-1 741-975 R | AGCTTCGCATACACGACCAA |
